# ODFormer: a Virtual Organoid for Predicting Personalized Therapeutic Responses in Pancreatic Cancer

**DOI:** 10.1101/2025.07.08.663664

**Authors:** Jing Xu, Xiaolu Yang, Yunguang Li, Huan Wang, Yikai Li, Shijie Tang, Shiwei Guo, Jiawei Zou, Xiaohan Shi, Zexi Chen, Chaoliang Zhong, Penghao Li, Wei Jing, Kailian Zheng, Xinqian Wu, Dong Gao, Luonan Chen, Gang Jin

## Abstract

Pancreatic cancer (PC) patient-derived organoids (PDOs) faithfully recapitulate therapeutic responses but face clinical translation barriers, including high costs and technical complexity. To address these problems and the lack of frameworks for PDO-based drug-response assays, we developed ODFormer, a computational framework that simulates PC PDOs to predict clinically actionable, patient-specific drug responses by integrating transcriptomic and mutational profiles. ODFormer first employed two encoders, pretrained on 30,000 pan-cancer bulk transcriptomics and 1 million PC single-cell profiles respectively, to distil tissue-and organoid-specific representations. Then, trained on our curated 14,000 PDO drug-response assay (across 183 PDOs and 98 drugs) using a transformer–augmented hybrid contrastive network, ODFormer significantly outperformed state-of-the-art methods, notably achieving a PCC >0.9 in predicting standardized drug response. Multi-cohort retrospective analyses further demonstrated that ODFormer-guided personalized therapy significantly improves clinical outcomes, without requiring physical organoid assays. Furthermore, ODFormer identified novel clinico-biological PC subtypes and revealed therapy resistance biomarkers by stratifying predicted responders and non-responders. These were validated using independent datasets including TCGA-PDAC. Notably, ODFormer-guided treatment efficacy showed high concordance with prospective clinical responses by CA19-9.

## Introduction

Personalized drug therapy selection plays a crucial role in optimizing clinical outcomes for cancer patients, particularly those with pancreatic cancer (PC), due to its aggressive nature and poor prognosis ^1–4^. To tailor therapy more precisely, a spectrum of preclinical models has been deployed. Historically, two-dimensional (2D) cell-line monolayers have served as the workhorse for high-throughput drug screening, but their flat architecture fails to capture essential tumor-stroma crosstalk, three-dimensional diffusion gradients, and heterotypic cell–matrix interactions, limitations that significantly curtail their ability to predict in vivo efficacy^5, 6^. In contrast, PDOs systems preserve original tumor architecture through 3D extracellular matrix embedding while maintaining autologous stromal components like cancer-associated fibroblasts and immune cells^7^. This biomimetic reconstruction enables more clinically relevant drug response profiling, with validation studies demonstrating high concordance between PDOs predictions and patient responses across multiple malignancies, including colorectal cancer^8^, pancreatic ductal adenocarcinoma^9, 10^, ovarian carcinoma^11^, and glioblastoma^12^. Nonetheless, both conventional 2D cultures and more physiologically relevant 3D models, such as PDOs, remain constrained by lengthy establishment timescales (often several weeks) and high per-sample costs. Notably, PDO generation typically necessitates substantially longer timelines, frequently exceeding the critical window for treatment selection in advanced-stage malignancies, while incurring costs of thousands of dollars per screen. These limitations render routine clinical implementation currently impractical. Against these backdrop, computational modeling emerges as a time-and resource-efficient adjunct: in silico approaches have demonstrated potential to predict drug response, thereby offering a scalable route to accelerate drug prioritization and combination screening^13–17^.

The development of accurate computational predictors for drug response critically depends on the availability of large, well-annotated pharmacogenomic datasets for both model training and validation. To date, two-dimensional cancer cell lines have dominated this field owing to their ease of culture and highly standardized protocols, which together have enabled the assembly of resources such as the Cancer Cell Line Encyclopedia (CCLE)^5^ and the Genomics of Drug Sensitivity in Cancer v1 and v2 (GDSC)^18^ and Cancer Therapeutics Response Portal (CTRP)^19, 20^. Leveraging these repositories, early computational frameworks, ranging from machine-leaning based regression models to deep-learning based regression models^14^, demonstrated promising performance under cross-validation and, in some cases, were confirmed by targeted cell lines^21, 22^. Despite these achievements, three interrelated challenges have emerged. First, most existing models are trained on cell line–drug response datasets, which lack the physiological relevance. As a result, predictions based on homogeneous monolayer cultures often fail to capture the architectural and molecular complexity of in vivo tumors, leading to poor concordance with patient outcomes and limited clinical applicability. Second, the majority of current models have not been rigorously validated using real-world clinical datasets—particularly those reflecting patient-specific endpoints—thereby restricting their utility in precision oncology. Third, conventional transcriptomic feature extraction methods often overlook biologically meaningful, dataset-specific information. These approaches typically reduce dimensionality based solely on mathematical criteria, without preserving latent biological signals, which undermines their interpretability and downstream utility. At the same time, there is a pressing need for AI models specifically designed for organoid pharmacogenomics^23^. With the anticipated growth of public omics datasets derived from PDOs, it is critical to develop computational frameworks tailored to the unique characteristics of organoid data^24, 25^. They not only enable the establishment of benchmarking frameworks and facilitate mechanistic insights into drug response, but also provide a robust foundation for accurately predicting patient-specific responses in translational contexts.

To address these critical limitations, we developed ODFormer (**O**rganoid-Derived **D**rug-response Trans**former**), an AI-driven "virtual organoid" platform designed to predict PC patient-specific drug responses by integrating tumor transcriptomes, mutation profiles, and drug chemical structures. This new approach eliminates the need for generating PDOs in vitro or conducting additional PDOs drug screening experiments, thereby overcoming key bottlenecks in personalized drug testing. Our approach is underpinned by three synergistic advances. First we developed a foundation-model-style pretraining paradigm, inspired by large-scale pretraining models shown to greatly enhance model generalizability across downstream tasks in single-cell and beyond^26^ ^27, 28^, to distil tissue- and organoid-specific representations. Second, we assembled the largest compendium to date of drug response profiles in PC PDOs, comprising over 14,000 dose–response area under the curve (DR-AUC) measurements across 183 PDO lines and 98 therapeutic agents^10, 29^. These serve as foundational resources for training and evaluating predictive models. Third, we developed a transformer-based deep learning framework that jointly encodes multi-omics patient profiles and drug molecular descriptors. This architecture is optimized using a hybrid contrastive-loss strategy, which adaptively calibrates drug-specific and response-specific thresholds to balance triplet-based relational constraints with global embedding consistency, thereby enhancing the fidelity of patient–drug interaction representations.

ODFormer demonstrates robust predictive power across diverse evaluation paradigms, significantly surpassing current state-of-the-art approaches. In particular, it achieves a Pearson correlation coefficient (PCC) greater than 0.90 for standardized drug-response prediction. This high accuracy holds true whether assessed at the single-drug or combination-therapy or drug-sensitivity task for an individual-patient, as confirmed by internal cross-validation and rigorous zero-shot evaluations on held-out cohorts. Multi-cohort retrospective analyses reveal its clinical translational potential: ODFormer-guided personalized therapy selection shows significant improvement in patient outcomes while reducing dependence on physical organoid screening. Beyond therapeutic prediction, the model identifies four novel PC subtypes with distinct molecular signatures and differential drug response patterns. These findings were validated across independent external cohorts—including patient-derived xenograft (PDX) models, our own prospectively collected internal cohort, and the TCGA-PDAC dataset—thereby demonstrating its broad applicability and robustness. Mechanistic interrogation of responder/non-responder predictions reveals resistance-associated genes, such as MUC2 and SPINK4. Importantly, prospective validation in 16 treatment-naïve patients demonstrates 68% accuracy in correlation with CA19-9 biomarker response. This multidimensional validation establishes ODFormer as a virtual organoid that not only suggests effective PC patient-specific therapy but also simultaneously advances both computational oncology and biological understanding.

## Results

### The overview of ODFormer

To address the substantial cost and time demands of generating PDOs and performing drug-sensitivity assays, we developed ODFormer, a “virtual organoid” framework to predict personalized drug DR-AUC values directly from tumor transcriptomic and mutational profiles. ODFormer comprises three interlinked components: data integration, model architecture, and clinical translation (Fig. 1).

**Figure 1.**
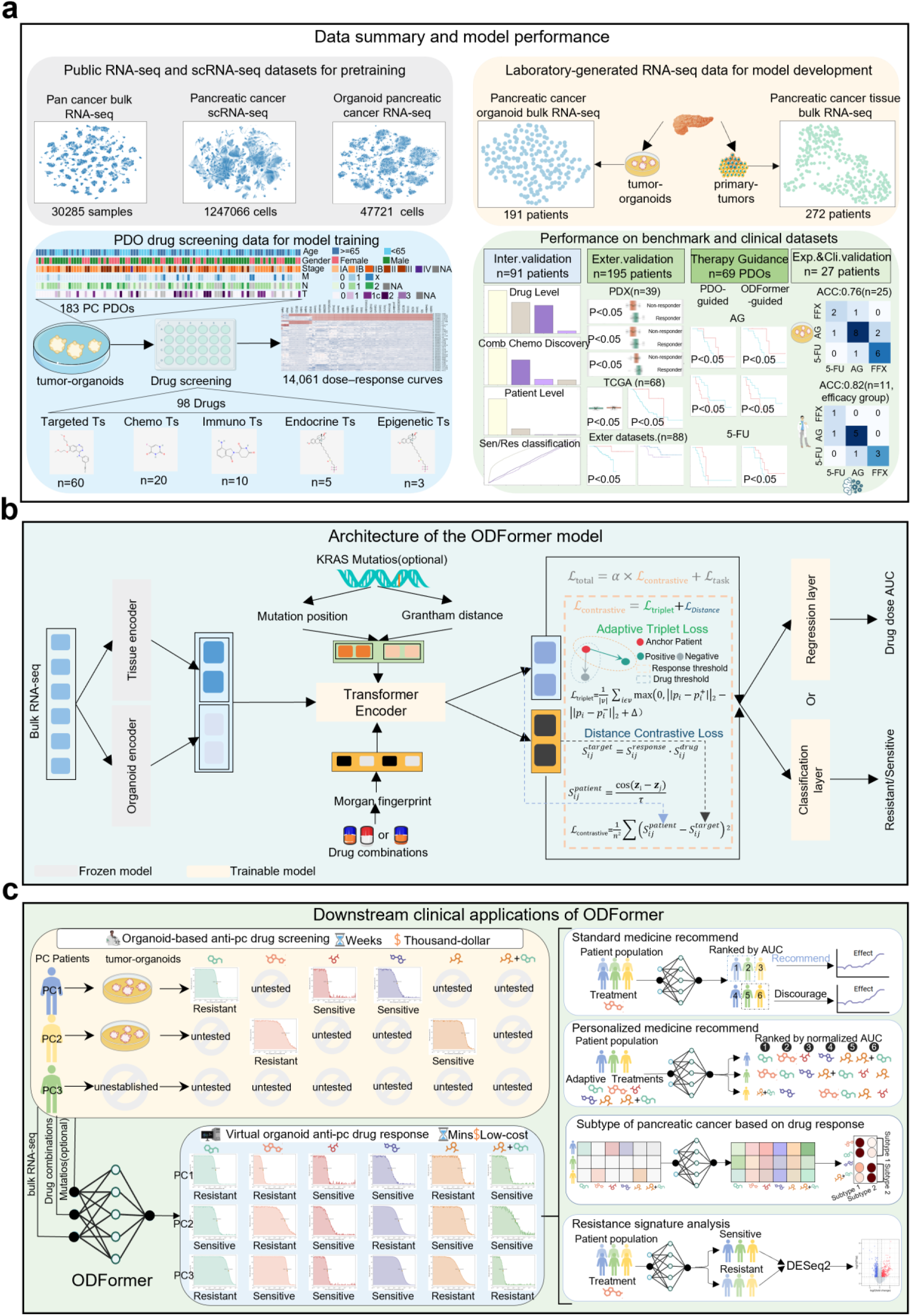
Overview of ODFormer framework and its clinical applications. (**a**) Data Overview: Overview of data resources and model development framework. Public RNA-seq (30,285 samples) and scRNA-seq (1,247,066 cells) datasets were used for model pretraining, while laboratory-generated RNA-seq data (335 pancreatic cancer samples) and patient-derived organoid (PDO) drug screening data (14,061 dose-response curves across 183 PDOs and 98 drugs) formed the training and evaluation datasets. The ODFormer model was developed and validated through internal (n=91 patients), external (n=82 patients), PDX models (n=39 samples), TCGA-PDAC (n=68 patients) and experimental (n=24 PDOs), and clinical (n=16 patients) cohorts. (**b**) Model Architecture: Architecture of the ODFormer model for personalized drug recommendation. The framework integrates tissue bulk RNA-seq data processed through a Transformer encoder and drug molecular features via Morgan fingerprints. The model simultaneously predicts drug sensitivity (resistant/sensitive classification) and dose-response AUC values (DR-AUC). KRAS mutation information is optionally incorporated using Grantham distance metrics. (**c**) Clinical Applications: Clinical implementation workflow and applications of ODFormer. The model imputes (DR-AUC) for patients and enables four key clinical utilities: (1) General therapeutic recommendations using tumor organoid drug screening; (2) Personalized treatment prioritization ranked by normalized AUC; (3) Adaptive treatment optimization combining multiple therapeutic strategies; (4) Pancreatic cancer subtyping based on drug response patterns through resistance signature analysis.

To establish foundational biological representations, we collected a diverse compendium of RNA-seq data comprising 30,000 pan-cancer bulk RNA-seq samples, 1 million scRNA-seq data and 47,721 organoid single cells from TISCH2^30^, GEO and TCGA^31^ to pretrained two encoders to learn robust biological representations (Fig. 1a). Next, we assembled the largest PC PDO drug-response dataset to date, comprising over 14,000 DR-AUC measurements across 183 PDOs treated with 98 therapeutic agents. In parallel, we generated laboratory datasets of 191 bulk RNA-seq profiles from pancreatic cancer organoids and 272 bulk RNA-seq profiles from matched primary tumor tissues. Among these 272 tumor samples, three cohorts were defined: (1) an internal validation cohort of 91 samples with matched PDO models and corresponding mutational profiles, (2) an external validation cohort of 82 samples lacking corresponding PDO data, and (3) a prospective cohort of 27 samples with matched PDOs subjected to in vitro sensitivity assays for 5-fluorouracil (5-FU), Gemcitabine plus nab-Paclitaxel (AG), and FOLFIRINOX (FFX), of which 16 patients had available clinical treatment records. These datasets enable rigorous internal and external validation, experimental verification, and preliminary clinical testing of ODFormer predictions.

ODFormer employs a transformer-based dual-encoder architecture specifically designed to address domain shifts between in vitro organoids and primary tumor tissues (Fig. 1b). This framework utilizes two distinct pre-trained transformer encoders: an Organoid encoder and a Tissue encoder (Supplementary Fig. 1). The Organoid encoder, pre-trained on a large compendium of organoid RNA-seq data using masked language modeling, captures the subtle, ex vivo expression signatures characteristic. Conversely, the Tissue encoder was pre-trained on pan-cancer data, optimizing its feature extraction for the intrinsic heterogeneity of patient samples. During processing, each transcriptomic profile from our clinical-omics cohort (191 PDOs and 272 tumor RNA-seq samples) is segmented into fixed-length vectors and independently encoded by both encoders, generating paired 128-dimensional embeddings. These organoid-derived and tumor-specific embeddings are concatenated into a unified feature vector. To integrate mutation information and drug information, we adopt the transformer architecture^32^, enabling the joint encoding of patient omics profiles and molecular drug features. Specially, we introduce a hybrid contrastive learning framework that synergistically integrates an adaptive triplet loss—featuring dynamic thresholds for drug similarity and response concordance—with a distance-based contrastive objective aligning patient embeddings to pharmacological and phenotypic profiles. This design enables the model to flexibly adjust positive and negative sample pairs, addressing data sparsity and heterogeneity inherent in organoid-based drug response studies. Our approach enhances the biological relevance of latent representations, offering a robust computational tool for precision medicine applications in computational biology and organoid research. Finally, ODFormer utilizes dual-task prediction heads that optimize continuous DR-AUC regression or binary sensitive/resistant classification for comprehensive drug response prediction.

The model addresses critical limitations of conventional organoid screening—such as time-intensive workflows (weeks) and high costs—by leveraging bulk RNA-seq and optional mutational data to impute/predict missing drug response profiles (DR-AUC values) in minutes. Key clinical utilities include: (1) General therapeutic recommendations derived from in vitro organoid drug sensitivity testing, bridging gaps in untested drug combinations; (2) Personalized treatment prioritization, where adaptive therapies are ranked by normalized AUC to optimize patient-specific efficacy; (3) Dynamic treatment optimization through combinatorial therapeutic strategies, enhancing precision medicine paradigms; and (4) Molecular subtyping of PC based on drug response patterns, validated by resistance signature analysis (e.g., DESeq2-driven differential expression). By synergizing experimental organoid models with AI-driven prediction, ODFormer accelerates translational oncology workflows while reducing costs, offering a scalable platform for both population-level and patient-centric decision-making.

### ODFormer models clinically realistic drug- and patient-level drug response

Patient-derived organoids have transformed cancer research by accurately recapitulating the histopathological and genomic features of human tumors, offering a physiologically relevant platform for in vitro drug sensitivity profiling to inform personalized treatment strategies and accelerate precision oncology^6^. However, the clinical translation of PDO-based approaches remains limited by the time-, labor-, and cost-intensive nature of organoid generation and drug screening workflows. To overcome these barriers, we leverage pharmacogenomic response data from PC-PDOs as training and validation sets for predictive modeling. To rigorously evaluate the clinical predictive power of our computational framework, ODFormer, we benchmarked it against state-of-the-art drug response prediction models, including DeepCDR^33^, BANDRP^34^, and PANCDR^35^. We systematically assessed performance through Pearson correlation coefficient (PCC) analysis, first at the drug-compound level and subsequently at the patient level.

To evaluate drug-level generalization, we modeled the prediction of organoid responses to novel, previously unseen compounds. This task holds substantial translational relevance, as it can reduce experimental screening efforts and accelerate the identification of patient-tailored therapies. It also offers potential to forecast the activity of investigational drugs, thereby informing early-stage development. Specifically, we utilized a comprehensive pharmacogenomic dataset in which each patient-derived organoid had been profiled against all 14 clinically approved pancreatic cancer drugs. We then performed five-fold cross-validation over the drug dimension, systematically withholding subsets of drugs for evaluation while training on the remaining compounds across all organoid lines. As illustrated in Fig. 2a, ODFormer demonstrated superior performance, achieving the highest PCC of over 0.7, significantly outperforming the second-best method by ΔPCC = 0.25. Next, we examined model performance for each individual drug. ODFormer consistently surpassed the second-best model (BANDRP), yielding the highest PCC across all drug predictions and patients predictions (Fig. 2b, Supplementary Fig. 2-3). Notably, this advantage was most pronounced for the clinically frontline combination regimens AG (PCC = 0.84, ΔPCC = 0.10 vs. BANDRP).

**Figure 2.**
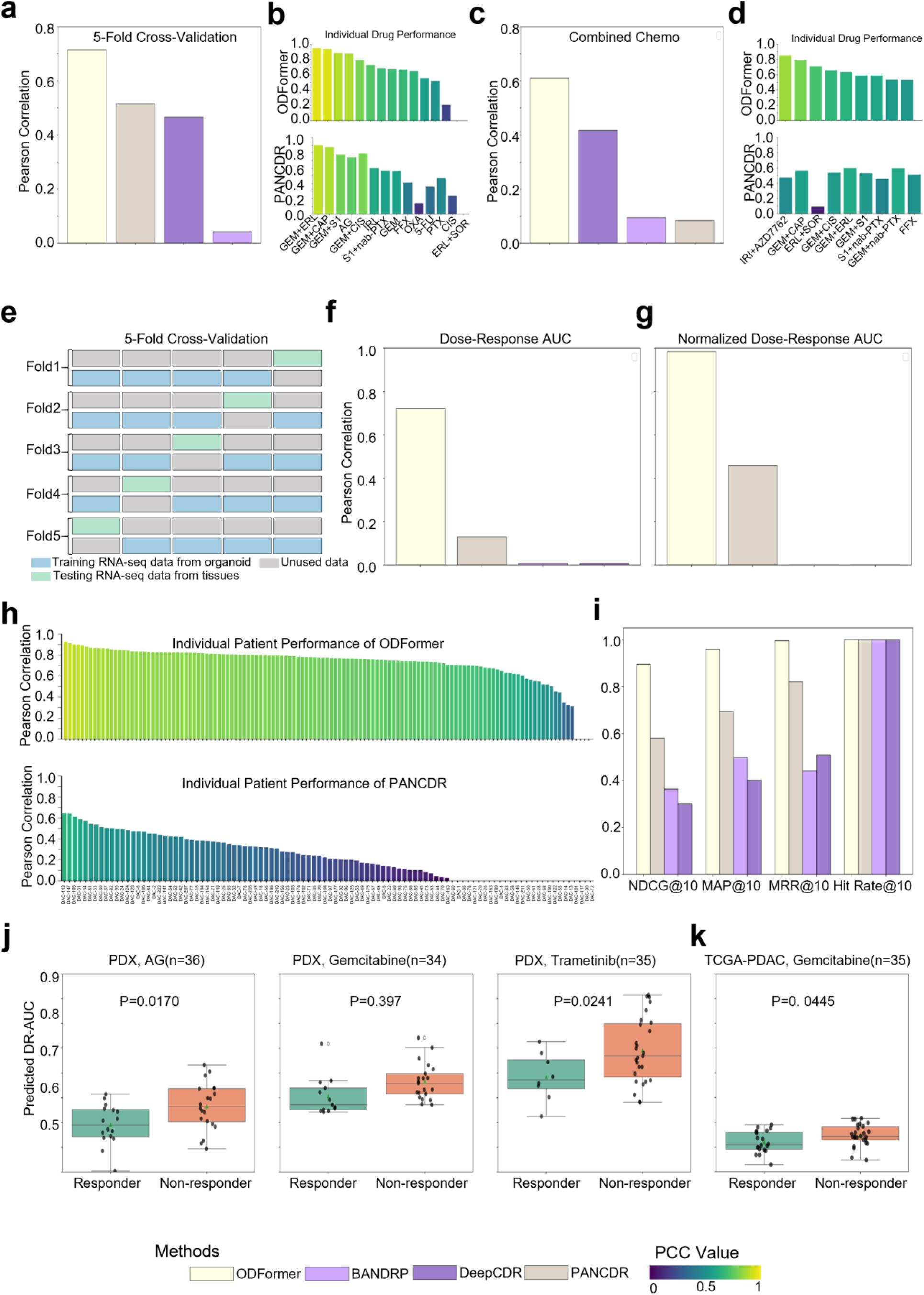
Evaluation of ODFormer for Drug Sensitivity Prediction and Survival Outcome Association in Pancreatic Cancer. (**a**) Bar plots comparing the Pearson correlation coefficient (PCC) achieved by ODFormer, DeepCDR, BANDRP, and PANCDR in predicting organoid responses to novel compounds using five-fold cross-validation. (**b**) Bar plots showing the PCC achieved by each model for individual drugs of ODFormer(top), PANCDR(bottom). (**c**) Bar plots comparing the PCC achieved by each model in predicting sensitivity to novel drug combinations using monotherapy data. (**d**) Bar plots showing the PCC achieved by each model for individual drugs within the predicted combinations of ODFormer(top), PANCDR(bottom). (**e**) Schematic illustrating the five-fold cross-validation strategy used to evaluate ODFormer’s performance in predicting drug responses directly from patient tumor tissue transcriptomes. (**f**) Bar plots comparing the PCC achieved by each model for DR-AUC prediction using patient tumor tissue transcriptomes. (**g**) Bar plots comparing the PCC achieved by each model for normalized DR-AUC prediction using patient tumor tissue transcriptomes. (**h**) Bar plots showing the individual patient performance of ODFormer and PANCDR, as measured by PCC. (**i**) Bar plots comparing the mean average precision (MAP), normalized discounted cumulative gain (NDCG), mean reciprocal rank (MRR), and hit rate achieved by each model in prioritizing effective therapies. (**j,k**) Box plots comparing the predicted DR-AUC values between treatment responders and non-responders in (**j**) patient-derived xenograft (PDX) models for AG, gemcitabine and Trametinib treatment arms and in (**k**)TCGA-PDAC for gemcitabine. P-values were calculated using the Mann-Whitney U test.

We next addressed the more challenging and clinically pertinent scenario of predicting responses to novel drug combinations using only monotherapy data. This is critical because combination therapies are frequently employed in oncology to enhance efficacy and combat resistance^36^. Here, we trained models using monotherapy DR-AUC data for 9 compounds across a cohort of organoids. We then evaluated their ability to predict sensitivity to 13 distinct, clinically relevant drug combinations within an independent validation set comprising 91 organoids. As shown in Fig. 2c, ODFormer maintained strong predictive capability for combination therapies, achieving a leading PCC above 0.6. Fig. 2d details performance for individual drugs within the predicted combinations (Supplementary Fig. 5-6). Crucially, ODFormer demonstrated superior performance in this zero-shot combination prediction setting. While PANCDR remained effective, its accuracy was generally lower than ODFormer’s, particularly for specific drug combinations.

To evaluate the model’s utility in scenarios where patient-derived organoids are unavailable, we simulated the generation of virtual organoids and predicted drug responses directly from patient tumor tissue transcriptomes. To rigorously assess ODFormer’s performance under clinically realistic and data-scarce conditions, we implemented a five-fold cross-validation strategy. In each fold, patient samples were divided into mutually exclusive training and testing cohorts to ensure no data leakage. Specifically, ODFormer was trained on organoid RNA-seq profiles paired with drug response data (light blue), and evaluated on unseen patient tissue RNA-seq–drug pairs (green). This strict cohort separation ensures that the test set comprises entirely novel patient samples, providing a robust estimate of the model’s generalizability in real-world clinical settings (Fig. 2e). ODFormer consistently outperformed existing approaches in drug response prediction. For DR-AUC, ODFormer achieved a 50% improvement in PCC over the second-best method (Fig. 2f). Similarly, for normalized DR-AUC prediction, ODFormer showed a 53% higher correlation (Fig. 2g). At the individual patient level, ODFormer was the only model to maintain a PCC exceeding 0.80 between predicted and observed responses across all cases, whereas competing models demonstrated notably inferior and less stable performance (Fig. 2h, Supplementary Fig. 7-8). While accurate response prediction is essential, clinical applicability ultimately depends on the model’s ability to prioritize effective therapies. To evaluate this, we assessed ODFormer’s ranking performance using four established metrics: mean average precision (MAP), normalized discounted cumulative gain (NDCG), mean reciprocal rank (MRR), and hit rate (Methods). ODFormer exhibited consistently strong performance across all ranking criteria, highlighting its potential for personalized drug prioritization in clinical practice (Fig. 2i).

To assess the translational relevance of ODFormer, we evaluated its performance in patient-derived xenograft (PDX) models and TCGA-PDAC cohorts. In PDX models, ODFormer significantly distinguished responders from non-responders, yielding lower predicted DR-AUC values in responders treated with AG (n = 36, P = 0.0170), gemcitabine (n = 34, P = 0.0397), and trametinib (n = 35, P = 0.0241) (Fig. 2j). Moreover, in TCGA-PDAC patients treated with gemcitabine (n = 35), ODFormer also identified response groups with significantly lower predicted DR-AUCs in responders (Fig. 2k, P = 0.0445).

The consistent predictive accuracy across different modelsand treatment regimens highlights ODFormer’s superior translational capability compared to conventional methods. Our systematic evaluations confirm the model’s robustness in predicting drug responses for novel patient samples, a critical requirement for clinical applications in precision oncology.

### ODFormer-guided Personalized Treatment Recommendations Improves Patient

#### Overall Survival

Having validated ODFormer’s predictive performance through internal cross-validation, we next assessed its potential as a virtual organoid for precision oncology using one inner pancreatic cancer cohorts and two independent pancreatic cancer cohorts with available survival data. The inner cohort included 69 patients with matched clinical outcomes and PDO drug-screening data. The first independent cohorts comprised 82 patients with clinical outcomes but without matched PDO data. The second independent cohorts included 65 patients from TCGA-PDAC.

In standard population-based organoid drug screening protocols, individual drugs are tested on organoids derived from multiple patients. Based on their DR-AUC values (ranked from low to high), patients were stratified into three groups: the top 30% were defined as the sensitive (sen) group, and the bottom 30% as the resistant (res) group. Theoretically, patients in the sensitive group who received the recommended drug should exhibit significantly longer survival compared to those who did not, whereas patients in the resistant group should show significantly shorter survival if treated with the ineffective drug. To evaluate this hypothesis, we first analyzed DR-AUC values derived from real PDO drug sensitivity experiments, and compared survival outcomes in both sensitive and resistant groups, according to whether patients received PDO-guided treatment recommendations. To validate the predictive utility of ODFormer, we then repeated the above analysis using DR-AUC values predicted by ODFormer. Specifically, we applied population-based drug recommendations guided by ODFormer to stratify patients into sensitive and resistant groups, and compared their survival outcomes with those stratified using real PDO-derived DR-AUC. This allowed us to assess whether ODFormer-guided treatment recommendations align with theoretical expectations and mirror the results of PDO-guided clinical decision-making.

Since AG and 5-FU was commonly prescribed to the patients in our cohort, we focused on the two drugs for further validation. To evaluate the predictive accuracy of ODFormer for AG response, we first assessed its overall performance across PDO samples. ODFormer achieved a high PCC of 0. 0.845 between predicted and observed drug responses (Fig. 2b, Fig. 3a, top). Stratified analysis further revealed strong subgroup performance: in the top 30% of drug-sensitive patients, ODFormer achieved a PCC of 0.701 with a root mean square error (RMSE) of 0.115; in the bottom 30% drug-resistant group, the PCC increased to 0.771 with an RMSE of 0.067 (Fig. 3a, top and bottom), suggesting superior accuracy in both extremes of response.

**Figure 3.**
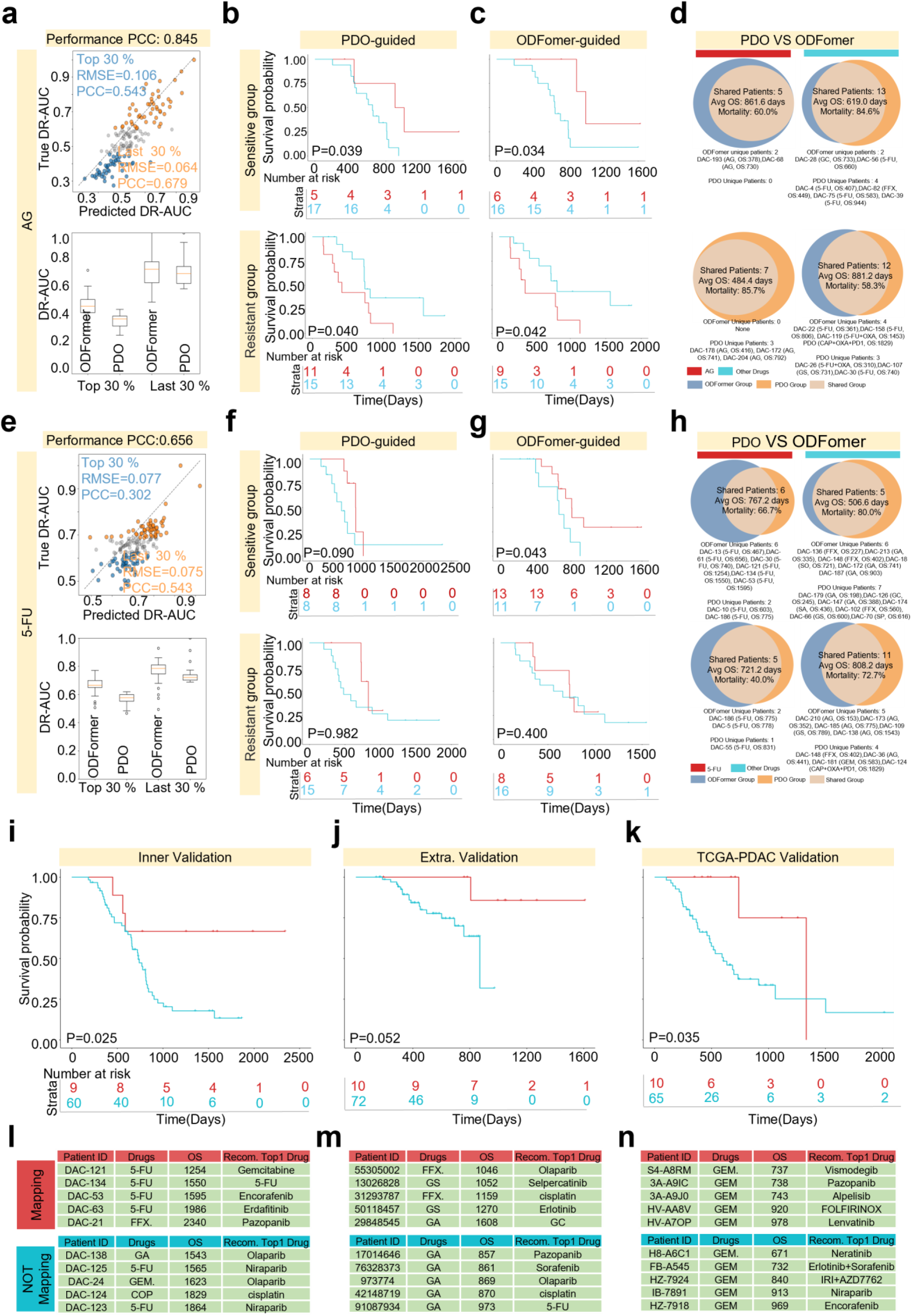
ODFormer-Guided Drug Sensitivity Prediction and Survival Outcomes in PDAC. (**a**) Scatter plots showing the correlation between predicted and observed DR-AUC values for AG in PDO samples. The top and bottom 30% of drug-sensitive and drug-resistant patients are highlighted, with corresponding PCC and RMSE values. Box plots display the distribution of DR-AUC values for ODFormer predictions and PDO measurements. (**b-c**) Kaplan-Meier survival curves comparing overall survival of patients treated with AG based on PDO-guided (**b**) and ODFormer-guided treatment recommendations in sensitive (top) and resistant (bottom) groups. P-values were calculated using the log-rank test. (**d**) Venn diagrams illustrating the overlap between patients classified as sensitive or resistant by PDO and ODFormer for AG. Median OS and mortality rates are shown for each group. (**e-g**) Similar analysis as in (a-c) for 5-FU. (**h**) Venn diagrams illustrating the overlap between patients classified as sensitive or resistant by PDO and ODFormer for 5-FU. Median OS and mortality rates are shown for each group. (**i-j**) Kaplan-Meier survival curves comparing OS of patients in internal (**i**) and external (**j**) cohorts, stratified by whether their actual treatment matched the top 10 drug recommendations predicted by ODFormer. P-values were calculated using the log-rank test.

To assess the clinical relevance of ODFormer predictions, we conducted survival analyses based on treatment recommendations guided by PDO-derived and ODFormer-predicted DR-AUC. In the PDO-guided group, AG-treated patients in the sensitive cohort showed significantly prolonged OS compared to those receiving alternative treatments (Fig. 3b, top; P = 0.039). Conversely, AG-treated patients in the resistant cohort exhibited a trend toward shorter OS (Fig. 3b, bottom; P = 0.040), aligning with in vitro DR-AUC estimates. Similarly, ODFormer-guided recommendations yielded consistent survival stratification: in the predicted-sensitive group, AG treatment was associated with significantly improved OS (Fig. 3c, top; P = 0.034), whereas in the predicted-resistant group, AG treatment resulted in significantly poorer survival outcomes (Fig. 3c, bottom; P = 0.042). To further examine concordance between the two approaches, Venn diagram analyses were performed. Among patients classified as sensitive by ODFormer (n = 13), the median OS was 619.0 days with a mortality rate of 84.6%; in contrast, ODFormer-predicted resistant patients (n = 12) had a shorter median OS of 581.2 days and a lower mortality rate of 58.3% (Fig. 3d; Supplementary Table 1). Despite partial overlap with PDO-based classifications, ODFormer provided complementary patient stratification, offering a data-driven and potentially more clinically informative tool for guiding chemotherapy decisions (other methods performance seen in Supplementary Fig. 9-11).

We next assessed the generalizability of ODFormer to other chemotherapeutic agents by evaluating its performance in predicting responses to 5-FU, a cornerstone drug in PDAC treatment. Although the overall predictive accuracy for 5-FU was lower than that observed for AG, ODFormer still achieved a moderate PCC of 0.656 (Fig. 3e, top), indicating reasonable predictive capacity. In the top 30% of patients classified as most sensitive, ODFormer achieved a PCC of 0.302 (RMSE = 0.077), while in the bottom 30% most resistant patients, predictive accuracy improved substantially, with a PCC of 0.543 (RMSE = 0.075) (Fig. 3e). Notably, survival analysis showed that ODFormer-guided treatment decisions in the sensitive subgroup yielded overall survival outcomes comparable to those seen with AG (Fig. 3f, top). In contrast, neither PDO- nor ODFormer-guided strategies produced significant survival differences among patients in the resistant group (Fig. 3f–3g, bottom), consistent with the attenuated predictive signal in this cohort. These suggest that a simplistic classification based solely on the top or bottom 30% of DR-AUC values may not fully capture the complexity of real-world treatment response, underscoring the need for more nuanced stratification strategies in clinical applications. Venn diagram analyses of patient classification consistency revealed limited overlap between PDO and ODFormer models. Among the shared sensitive patients (n = 5), the median OS reached 506.8 days with an 80.0% mortality rate (Fig. 3h, top right; Supplementary Table 2), while shared resistant patients (n = 5) exhibited a median OS of 721.2 days and a mortality rate of 40.0% (Fig. 3h, bottom left; Supplementary Table 2). Interestingly, ODFormer-unique resistant patients (n = 11) showed prolonged median OS (808.2 days) and relatively low mortality (72.7%), suggesting potential underestimation of sensitivity by ODFormer in some cases (other methods performance seen in Supplementary Fig. 12-14).

To further assess the clinical utility of ODFormer, we ranked the top 10 drug recommendations for each patient based on 45 drugs predicted norm DR-AUC values. Patients were classified as matched if any of their top 10 ODFormer-predicted drugs aligned with their actual treatment regimen; otherwise, they were considered non-matched. Survival analyses across three pancreatic cancer cohorts—an internal cohort (n = 69), an external real-world cohort (n = 82), and TCGA-PDAC (n = 65)—consistently showed improved outcomes for the matched group. In the internal cohort, matched patients exhibited significantly prolonged overall survival (OS) compared to non-matched patients (Fig. 3i; Supplementary Table 3; P = 0.025). This trend was recapitulated in the external cohort (P = 0.052; Fig. 3j) and the TCGA-PDAC cohort (P = 0.035; Fig. 3k; Supplementary Table 4), underscoring the model’s robustness across heterogeneous datasets. Collectively, these results demonstrate the potential of ODFormer to inform personalized treatment strategies and improve clinical outcomes. Comparative results from baseline models are shown in Supplementary Fig. 16.

These demonstrate that ODFormer effectively replicates the drug sensitivity prediction performance of PDO-based assays with comparable accuracy. The high concordance between ODFormer predictions and PDO-derived results further supports its reliability in guiding drug selection for pancreatic cancer. Its combination of accuracy, clinical relevance, and operational efficiency positions ODFormer as a promising tool for personalized cancer therapy, potentially reducing the dependency on resource-intensive experimental platforms.

### Imputed Drug Response Reveals Novel Therapeutic Subtypes

One of the key advantages of organoid-based models lies in their accessibility and scalability, which facilitate comprehensive molecular profiling across multiple omics layers—including transcriptomics and proteomics—to elucidate mechanisms underlying disease progression^37^. In pancreatic ductal adenocarcinoma, which exhibits high molecular and therapeutic heterogeneity, omics-based stratification has led to the identification of novel subtypes with distinct prognostic implications^38^. Nevertheless, drug response remains one of the most direct phenotypic readouts of tumor behavior. Despite its potential, systematic subtyping of PDAC based on PDOs pharmacological profiles remains unexplored, largely due to the scarcity and incompleteness of existing datasets. To address this gap, ODFormer leverages partially observed PDO-drug response data to impute missing values, thereby enabling subtype discovery grounded in pharmacological phenotypes.

In our study, among the 98 drugs tested, approximately 65% of patients exhibited missing response values, with individual-level sparsity ranging from 10% to 50% (Fig. 4a). At the drug level, no compound had complete coverage across all patients (Fig. 4b). To address this, we employed ODFormer to impute the missing DR-AUC values, enabling comprehensive analysis across the full PC PDOs-drug matrix. Overall, the imputed DR-AUC values showed strong concordance with the distribution of observed values (Fig. 4c). Representative examples at both the patient level (Fig. 4d) and drug level (Fig. 4e) demonstrate that, apart from cases with over 30% missingness, the imputed values closely resembled the empirical distributions.

**Figure 4.**
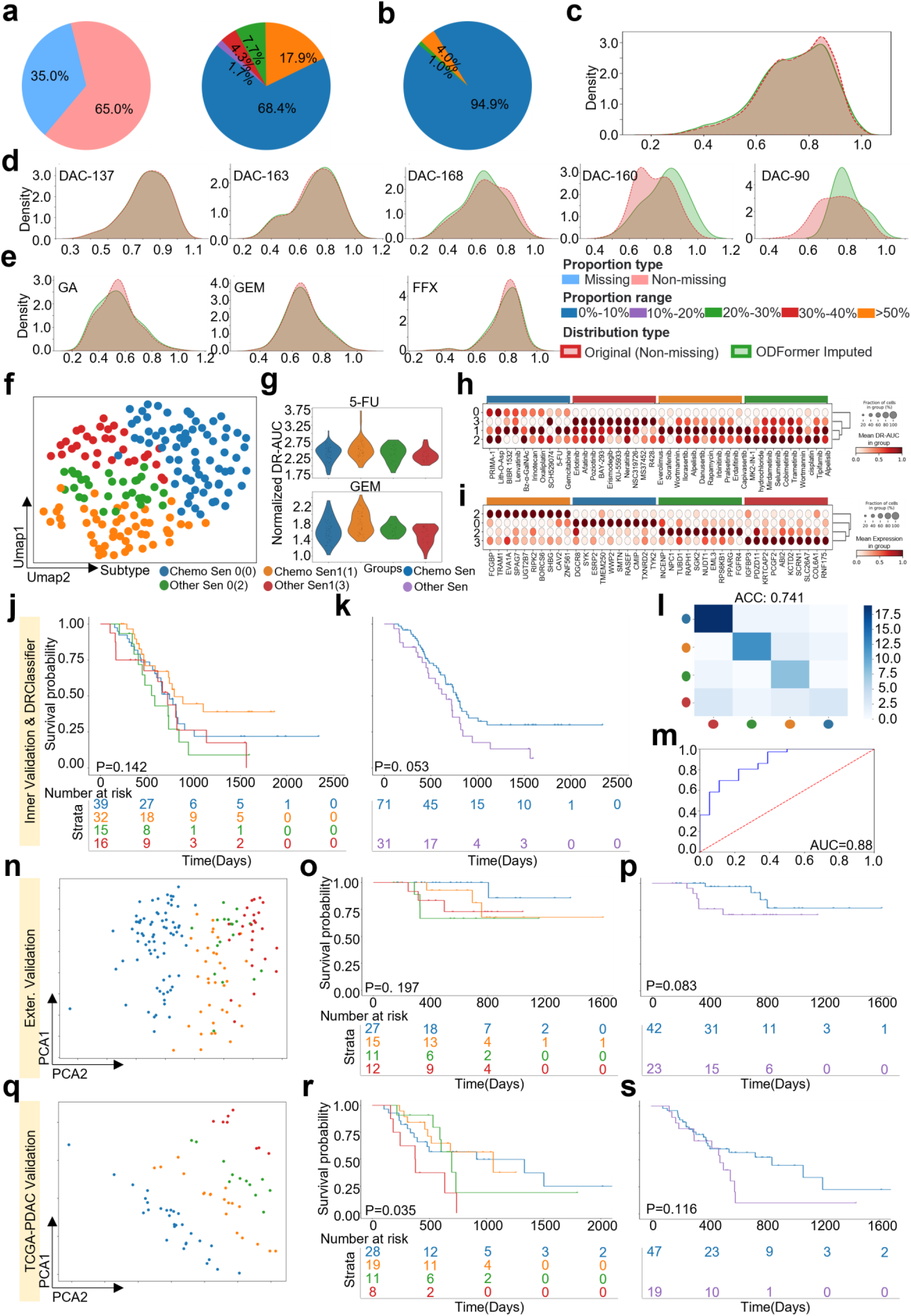
Drug Response Subtyping of PDAC Using ODFormer Imputation. (**a and b**) Pie charts showing the extent of missing drug response data across patients, with ∼65% of patients exhibiting missing response values. (**c**) A density plot comparing observed and imputed drug response area under the curve (DR-AUC) values, demonstrating strong concordance. (**d and e**) Examples of imputed DR-AUC distributions for individual patients and drugs, respectively. (**f**) Unsupervised clustering of patients using standardized imputed DR-AUC values reveals three distinct subgroups. (**g**) Drug sensitivity profiles of the identified subgroups, highlighting the third subgroup’s elevated sensitivity to chemotherapy agents such as 5-FU and Gemcitabine. (**h**) Marker drug identification for each subgroup, with the chemotherapy-sensitive group primarily exhibiting sensitivity to chemotherapy agents. (**i**) Transcriptomic signatures supporting the biological coherence of the subgroups, validated by Gene Ontology and Reactome pathway enrichment analyses. (**j and k**) Kaplan-Meier survival analysis for each subgroup, with the chemotherapy-sensitive subgroup showing significantly prolonged overall survival (log-rank p=0.05). (**l,m**) Machine learning classifier (DRClassifier) trained on imputed drug sensitivity data demonstrates robust performance on external datasets, with superior survival outcomes for the chemotherapy-sensitive subgroup. (**n-p**) Independent cohort validations with survival outcomes. (**q-s**) External validation in TCGA-PDAC cohort.

After standardizing the imputed DR-AUC values, where higher values indicated greater drug sensitivity, unsupervised clustering revealed four distinct patient subgroups (Fig. 4f). Drug response analysis showed that the first and second subgroup exhibited the highest sensitivity to both 5-FU and Gemcitabine, suggesting a distinct sensitivity profile (Fig. 4g). To further characterize each subgroup, we identified group-specific marker drugs (Fig. 4h). Most marker drugs in the first and second subgroups were conventional chemotherapeutic agents, prompting us to label them as Chemo Sen 0 (0) and Chemo Sen 1 (1), respectively. The remaining subgroups were termed Other Sen 0 (2) and Other Sen 1 (3), reflecting their distinct drug response profiles. This stratification was strongly supported by corresponding transcriptomic signatures (Fig. 4i), confirming the biological coherence of the defined subgroups. In particular, MSigDB Hallmark gene set enrichment analyses of Chemo Sen 0 (0) and Other Sen 0 (2) further validated the biological distinctiveness of these clusters^39, 40^ (Supplementary Figs. 16–18; Supplementary Tables 6–9).

Next, we also conduct survival analysis of the four subgroups using Kaplan-Meier curves revealed differences (Fig. 4j). Notably, the chemotherapy-sensitive subgroup, which received chemotherapy regimens, showed significantly prolonged overall survival compared to the other subgroups (Fig. 4k; P = 0. 0.053).

These results not only validate our imputation strategy but also reveal a clinically meaningful patient subgroup with increased sensitivity to standard chemotherapy. To assess the robustness of these drug-response subtypes, we trained a machine-learning model, DRClassifier, using 70% of the imputed drug sensitivity profiles from 180 samples. The classifier achieved 74% accuracy in the four-class classification task (Fig. 4l), and a binary classification of Chemo Sen versus all Other Sen yielded an area under the ROC curve of 0.88 (Fig. 4m).When evaluated on independent external cohorts with available predicted drug-response and survival data, DRClassifier maintained consistent performance: patients predicted as Chemo Sen 0 again showed significantly prolonged overall survival in both the external validation set and the TCGA-PDAC cohort(Fig. 4n-s).

The consistent survival advantage observed in Chemo Sen patients across multiple datasets highlights the clinical relevance of ODFormer, suggesting it as a novel, actionable subtype that can inform treatment selection in precision oncology. These underscore the utility of ODFormer in leveraging drug sensitivity predictions to achieve biologically and clinically meaningful patient stratification, offering an innovative framework for addressing the unmet need in pancreatic cancer subtyping.

### Resistance Biomarker Identification with ODFomer

In the previous section, we demonstrated the accuracy of the model in predicting DR-AUC values, which represent continuous outcomes. However, predicting a patient’s sensitivity to a specific drug is more effective and expedient when framed as a binary classification task. To this end, we adopted a binary classification framework by converting continuous DR-AUC values into discrete sensitivity labels (Fig. 5a, sensitive vs. non-sensitive, Methods). ODFormer exhibited superior performance, achieving a ROC AUC of 0.767 in stratifying PDAC patients (Fig. 5b), significantly outperforming both DeepCDR (AUC = 0.513) and PANCANR (AUC = 0.478). The model also maintained strong precision-recall characteristics (AUC = 0.810), a critical feature for clinical datasets with class imbalance, where minimizing false negatives is essential. A stability analysis further validated the robustness of ODFormer, with patient-specific performance ranging from AUC 0.72 to 1.00 (Fig. 5c).

**Figure 5.**
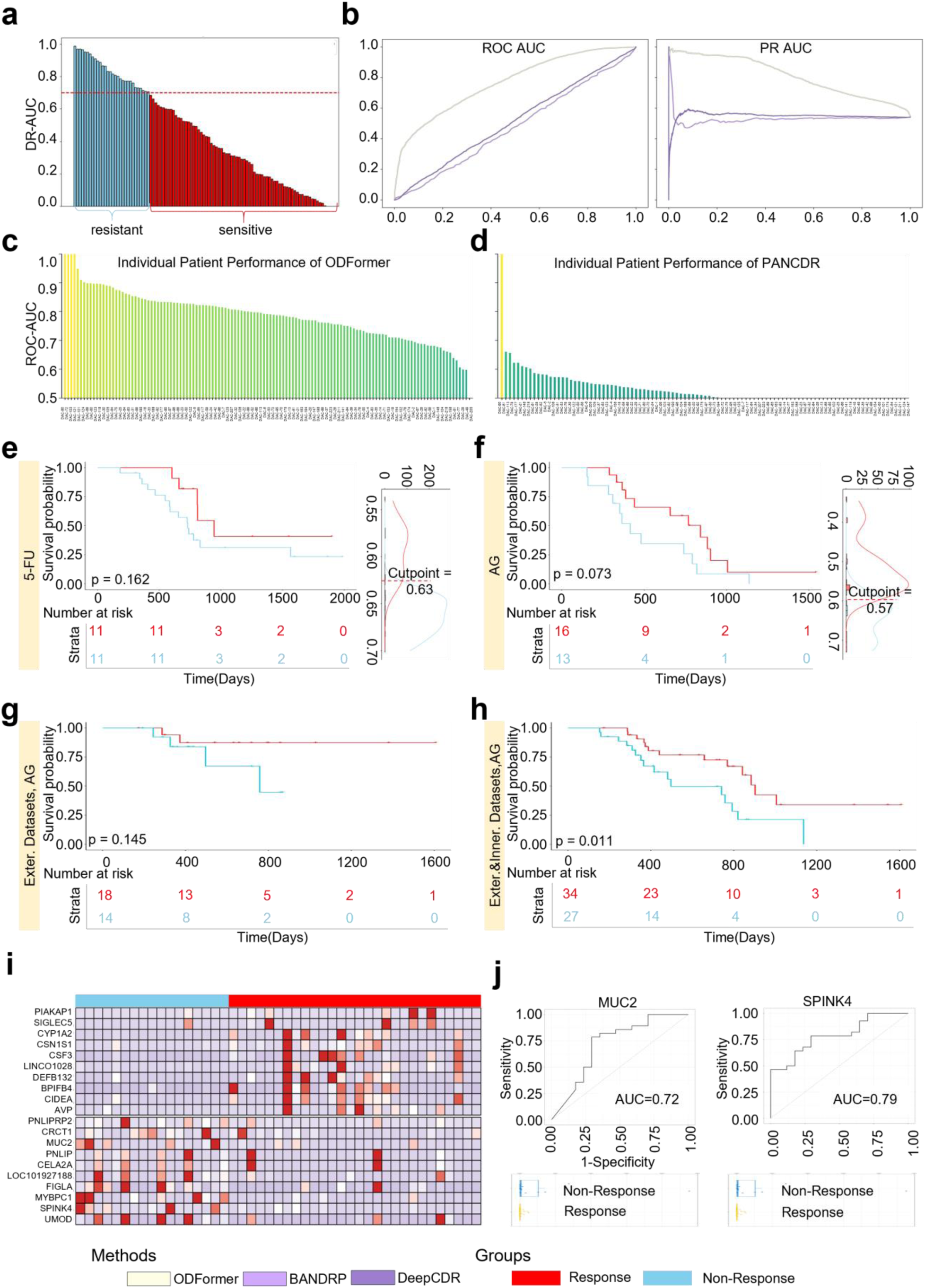
Evaluation of ODFormer for Drug Sensitivity Prediction and Survival Outcome Association in Pancreatic Cancer. (**a**) Schematic illustrating the conversion of continuous DR-AUC values into binary sensitivity labels (sensitive vs. non-sensitive). (**b**) Receiver operating characteristic (ROC) and precision-recall (PR) curves comparing the performance of ODFormer, DeepCDR, and PANCANR in predicting drug sensitivity. Area under the ROC curve (AUC) and area under the PR curve (AUPR) values are shown. (**c,d**) Waterfall plot showing the individual patient performance of ODFormer, as measured by AUC. Waterfall plot showing the individual patient performance of PANCANR, as measured by AUC. (**e-f**) Kaplan-Meier survival curves comparing overall survival (OS) of patients stratified based on 5-FU (**e**) and AG (**f**) sensitivity using OS-based cut-points. P-values were calculated using the log-rank test. Density plots show the distribution of DR-AUC values and the cut-point used for stratification. (**g**) Kaplan-Meier survival curves comparing overall survival of patients stratified based on AG sensitivity in the external cohort using survival-adapted labels. P-value was calculated using the log-rank test. (**h**) Heatmap showing differential gene expression between samples labeled by the external AG survival classifier and those from the internal cohort. Known AG-resistance markers (**i**)MUC2 and (**j**) SPINK4 are highlighted. Receiver operating characteristic (ROC) curves comparing the performance of ODFormer in predicting drug sensitivity in the internal cohort. Area under the ROC curve (AUC) values are shown.

Standardized binary drug-response measurements obtained through clinical trials remain the gold standard for evaluating therapeutic efficacy. Consequently, dichotomizing continuous DR-AUC values into “sensitive” versus “insensitive” categories typically relies on empirical cut-points (Fig. 3a–h) or purely statistical criteria, neither of which guarantee a direct link to true therapeutic efficacy. Overall survival, however, offers a more holistic, population-level proxy for treatment response, so we extended our framework by introducing survival-adapted labeling (Supplementary Note 1). We stratified 5-FU/AG–treated patients according to OS duration to define sensitive and insensitive groups. Although the OS differences between these two groups did not reach statistical significance, the observed trend aligned with our expectations (Fig. 5e,f). Leveraging these survival-adapted labels as training data, we then predicted AG sensitivity in an external cohort, achieving clear separation of OS curves (Fig. 5g). Finally, we performed differential transcriptomic analysis between samples labeled by the external AG survival classifier and those from our internal cohort, recovering known AG-resistance markers—MUC2^41^ and SPINK4^42^ (Fig. 5i,j; Supplementary Figure 19; Supplementary Table 10-11).

These establish ODFormer as a dual-capability platform, offering both drug response prediction and survival outcome association. By integrating pharmacogenomic data with clinical survival metrics, the platform addresses critical limitations in current therapeutic decision-making. Moreover, the concurrent identification of predictive biomarkers enhances its clinical utility by enabling mechanism-guided patient stratification, thereby advancing personalized oncology towards data-driven paradigms.

### Clinical Validation of ODFormer-Guided Personalized Therapy

To assess the clinical utility of the ODFormer framework, we conducted a prospective study involving 27 newly collected patient-derived samples, each paired with PDOs. Among these, 16 patients had complete clinical treatment records. ODFormer was employed to predict patient responses to commonly administered chemotherapy regimens, including 5-FU, AG, and FFX, with DR-AUC values serving as treatment recommendation scores (Methods). In parallel, drug sensitivity assays were performed on PDOs from all 27 patients to characterize resistance profiles against the same regimens. For 16 patients, clinical treatments were selected by physicians using standard clinical protocols and, where applicable, informed by PDO assay results.

As shown in Fig. 6a, patients in the concordant and effective group—those for whom both ODFormer recommendations and clinical prescriptions aligned—demonstrated significantly improved outcomes compared to patients in discordant groups. Notably, in cases P1–P9, ODFormer-predicted regimens were consistent with clinical decisions and were associated with substantial reductions in serum CA19-9 levels. In contrast, patients such as P10 and P11, whose treatments diverged from ODFormer predictions, experienced therapeutic failure, highlighting the model’s predictive robustness. While PDO-based guidance offered mechanistic insights into drug resistance—reflected in partial responses (e.g., moderate sensitivity in P1 and P4; resistance in P6, P8, and P13)—the variability in clinical outcomes suggested lower consistency compared to ODFormer.

**Figure 6.**
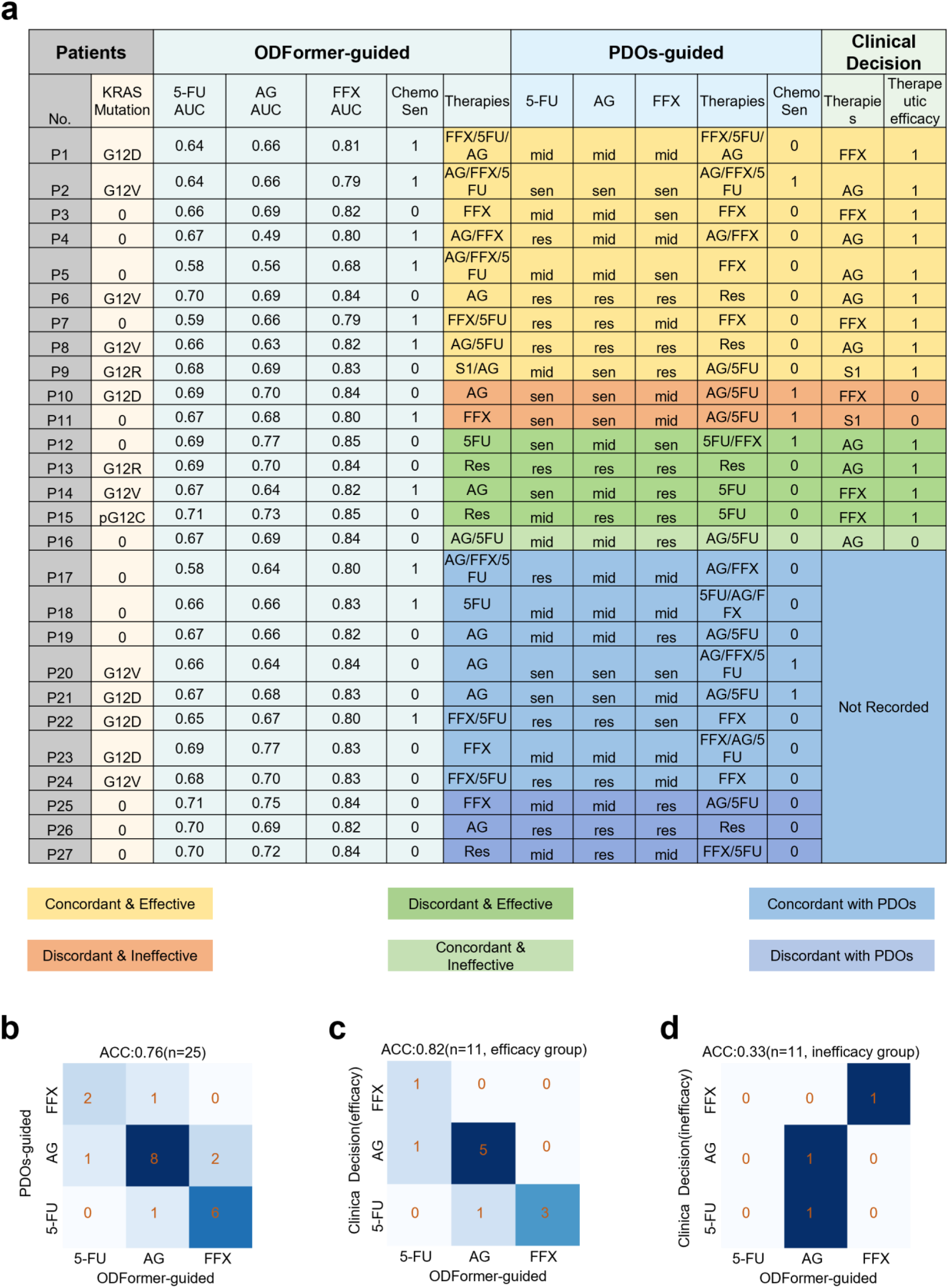
Concordance of ODFormer-Guided and PDO-Guided Treatment Recommendations with Clinical Outcomes in Colorectal Cancer Patients. (**a**) The table summarizes the predicted and observed responses to various chemotherapy regimens (5-FU, AG, FFX) in 24 patients with matched patient-derived organoids (PDOs). For each patient, KRAS mutation status, ODFormer-predicted AUC values for each drug, ODFormer-predicted chemotherapy sensitivity (Chemo Sen, binary), and PDO-predicted drug sensitivities (Sen = Sensitive, Mid = Intermediate, Res. = Resistant) are shown. The final column indicates the actual clinical treatment decision and therapeutic efficacy (1 = effective, 0 = ineffective). Rows are color-coded to indicate concordance between ODFormer prediction and clinical outcome (Concordant & Effective, Discordant & Effective, Discordant & Ineffective, Concordant & Ineffective), and concordance between PDO prediction and clinical outcome (Concordant with PDOs, Discordant with PDOs). (**b**) Confusion matrix comparing ODFormer and PDO-guided predictions (excluding patients recommended resistance by ODFormer); 76% overall concordance was observed., (**c**) ODFormer achieved 82% accuracy in matching clinical therapeutic outcomes. (**d**) Among patients who received clinically ineffective therapies, ODFormer matched physician decisions in only 33%, suggesting enhanced ability to reject non-beneficial treatments.

Overall, ODFormer-based recommendations showed a 76% concordance with PDO-based predictions across all patients, excluding cases where resistance was recommended. When benchmarked against actual clinical prescriptions, ODFormer matched the physician-selected regimens in 68.75% (11/16) of cases. Importantly, among patients who responded favorably to their treatments, ODFormer achieved a predictive accuracy of 82%. Conversely, in cases with clinically ineffective therapies, the model aligned with physician decisions in only 33%, underscoring its superior discriminative power in identifying efficacious treatment regimens.

In conclusion, this prospective evaluation supports the clinical relevance of ODFormer as a decision-support tool for personalized cancer therapy. By providing accurate predictions of therapeutic efficacy and aligning with real-world outcomes, ODFormer offers a promising framework to augment treatment planning and improve patient care in precision oncology.

## Discussion

Patient-derived organoids recapitulate tumor biology but are limited by slow generation, high cost, and sparse drug screening. Computational models offer promise but lack training and validation on PDOs with clinically relevant endpoints. To address these limitations, we developed ODFormer, a virtual organoid model that integrates transcriptomic, mutational, and drug structural data to predict patient-specific therapeutic responses in pancreatic cancer.

Compared to prior efforts largely reliant on monolayer cell-line pharmacogenomic datasets^43, 44^, ODFormer was trained and evaluated on a physiologically relevant dataset comprising our 14,000 organoid-derived drug response profiles from 183 patient-derived organoids across 98 compounds. This approach substantially alleviates a long-standing challenge in the field, namely the mismatch between in vitro model responses and in vivo clinical outcomes. By achieving a PCC exceeding 0.9 for standardized drug response prediction and consistently outperforming state-of-the-art models across diverse validation settings. This accuracy is particularly notable in zero-shot settings, such as drug combination prediction or patient tissue–based inference. These results support our hypothesis that biologically grounded, dual-stream pre-training, using 30,000 pan-cancer bulk transcriptomic profiles and 1 million prostate cancer single-cell profiles respectively, along with cross-attention-based fusion strategies, substantially improves the model’s generalizability. Furthermore, by employing contrastive learning objectives, ODFormer is capable of distinguishing subtle phenotypic heterogeneity among PDOs, a capability that is typically lacking in conventional regression-based models.

Importantly, ODFormer demonstrated the capacity to replicate and, in many instances, surpass the performance of physical PDO-based assays in stratifying patients for survival benefit. The clinical benefit of ODFormer-guided treatment selection was observed across multiple independent cohorts, strengthening the argument for its deployment in precision oncology pipelines. These suggests that our virtual organoid model may offer an alternative route to clinical decision-making, especially in scenarios where time or sample availability precludes conventional organoid generation. Perhaps most compellingly, our imputation of missing pharmacological data across a large organoid-drug matrix enabled the discovery of novel PDAC subtypes defined not by static molecular features, but by dynamic therapeutic responsiveness. These chemo-sensitive subtypes, validated and correlated with improved survival, underscore the potential of ODFormer not only as a predictive model, but as a hypothesis-generating tool. By shifting the focus from omics-based subtype classification to drug response-based stratification, our approach opens new avenues for therapeutic targeting and clinical trial design. Notably, our prospective validations, particularly the high concordance between ODFormer predictions and CA19-9 biomarker dynamics, further underscore its translational relevance.

Despite its strengths, ODFormer has certain limitations. First, its predictive accuracy for novel compounds or previously unseen patients is constrained by the chemical and biological diversity represented in the training dataset, highlighting the importance of continuous expansion and diversification of the pharmacogenomic corpus. Given the privacy concerns associated with patient data and the high cost of generating organoid-based datasets, we plan to explore federated learning^45, 46^ or swarm learning strategies^47, 48^ to enable decentralized model training without compromising data confidentiality. Second, although the current model incorporates transcriptomic and mutational data, the integration of additional omics layers—such as proteomics^49, 50^, single-cell transcriptomics^51, 52^, spatial transcriptomics^53^, and hematoxylin and eosin (H&E) histology images^54^—could further enhance predictive resolution^55^. Multimodal and multi-omics data offer complementary biological insights from different perspectives, which can improve both the generalizability and robustness of the model^56, 57^. However, these often suffer from missing modality issues, for which we plan to adopt large-model techniques to improve cross-modal stability and representation learning^28, 58, 59^. Third, while preliminary retrospective and prospective validations suggest potential clinical relevance of ODFormer, establishing its definitive utility will require randomized controlled trials (RCTs)^60^. Progress in this direction is currently limited by ethical considerations and the availability of large-scale, longitudinal datasets. Moving forward, we aim to further refine the model and expand clinical data collection to enable rigorous prospective evaluation. Expanding the application of ODFormer beyond pancreatic cancer is essential for realizing its broader potential as a pan-cancer predictive framework for therapeutic response. Currently, ODFormer has been evaluated primarily in the context of pancreatic ductal adenocarcinoma, where treatment options are largely limited to conventional chemotherapeutics. Future work will focus on adapting the model to other cancer types and incorporating a wider spectrum of therapeutic modalities, including targeted agents and immunotherapies^54, 61^. Additionally, interpretability of transformer-based architectures remains a technical frontier, requiring dedicated efforts to improve transparency and mechanistic insight extraction from deep learning models.

In summary, by enabling rapid and individualized therapeutic predictions and revealing previously unrecognized pharmacological subtypes, ODFormer not only bridges the gap between in vitro PDO assays and clinical application but also redefines the paradigm of drug response modeling. ODFormer constitutes both a conceptual and technical advancement in computational oncology. It provides a scalable, biologically informed, and clinically validated platform that circumvents the logistical limitations associated with physical organoid generation. As pharmacogenomic datasets continue to expand, ODFormer will be instrumental in realizing the full potential of precision oncology. Future developments may include the integration of additional omics layers, dynamic modeling of temporal drug responses, and adaptation to other tumor types. Although further clinical trials are necessary, particularly in first-line treatment settings, our findings support the adoption of virtual organoid frameworks as a foundational component of next-generation precision medicine.

## Methods

### Organoid Single-Cell Encoder

The Organoid Bulk Encoder is a deep learning model designed to process and encode high-dimensional transcriptomic data, initially pre-trained on PC scRNA-seq data and subsequently fine-tuned on organoid transcriptomic data. This two-stage training approach leverages the generalizable patterns learned from single-cell data to improve performance on organoid-specific tasks.

The pre-training phase begins with the preparation of scRNA-seq data, which is loaded and preprocessed using the scanpy library. The data is segmented into fixed-length vectors using a custom GeneEmbeding function, with a gap size of 128, to create a structured input representation. To simulate the sparsity and dropout effects commonly observed in single-cell data, 75% of the segments in each cell are randomly set to zero. The model architecture is based on a modified GPT (Generative Pre-trained Transformer) framework, incorporating a causal self-attention mechanism with ReLU activation instead of the traditional softmax. This modification enables the model to capture non-linear relationships in the data more effectively. The architecture includes multiple transformer blocks, each consisting of a multi-head self-attention layer and a feed-forward neural network, along with layer normalization and dropout layers to stabilize training and prevent overfitting. The model is pre-trained using the AdamW optimizer with a learning rate of 1e-4, a batch size of 32, and 100 epochs, minimizing the mean squared error (MSE) loss between predicted and actual gene expression profiles.

Following pre-training, the model is fine-tuned on organoid transcriptomic data to adapt it to the specific characteristics of organoid systems. The fine-tuning process uses the same architecture and optimization settings as the pre-training phase but focuses on organoid-specific datasets. This step allows the model to refine its learned representations, capturing organoid-specific patterns while retaining the generalizable features learned from single-cell data. The fine-tuned model generates embeddings that are highly relevant for downstream tasks such as clustering, visualization, and integration of organoid data. The final embeddings and model weights are saved for further analysis, enabling researchers to leverage the model for organoid-specific applications.

This two-stage training strategy—pre-training on single-cell data followed by fine-tuning on organoid data—ensures that the Organoid Bulk Encoder is both generalizable and highly specialized. By leveraging the rich information in single-cell datasets during pre-training, the model achieves robust performance on organoid data, making it a powerful tool for analyzing complex transcriptomic datasets in organoid systems.

### Tissue Bulk Encoder

The Tissue Bulk Encoder follows a similar training strategy to the Organoid Bulk Encoder but is specifically designed for bulk transcriptomic data. Instead of scRNA-seq data, the Tissue Bulk Encoder is pre-trained on pan-cancer bulk transcriptomic datasets, which encompass a wide range of cancer types and tissue contexts. This pre-training phase allows the model to learn generalizable patterns and features across diverse cancer types, capturing the heterogeneity and complexity inherent in bulk transcriptomic data. The architecture and training process mirror those of the Organoid Bulk Encoder, utilizing a modified GPT framework with a causal self-attention mechanism, ReLU activation, and transformer blocks. The model is trained to minimize the mean squared error (MSE) loss between predicted and actual gene expression profiles, with hyperparameters such as a learning rate of 1e-4, a batch size of 32, and 100 epochs.

After pre-training on pan-cancer bulk data, the Tissue Bulk Encoder can be fine-tuned on specific tissue or cancer types to adapt to the unique characteristics of the target dataset. This fine-tuning process enhances the model’s ability to capture tissue-specific or cancer-specific transcriptional patterns, making it highly effective for tasks such as cancer subtype classification, biomarker discovery, and integration of multi-omics data. By leveraging the broad knowledge gained from pan-cancer pre-training, the Tissue Bulk Encoder achieves robust performance on targeted datasets, providing a versatile and powerful tool for analyzing bulk transcriptomic data across a wide range of biological and clinical contexts. Together, the Organoid Bulk Encoder and Tissue Bulk Encoder demonstrate the flexibility and scalability of this training paradigm, enabling researchers to address diverse challenges in transcriptomics with tailored, high-performance models.

### Feature embeddings of RNA-seq/Driver-mutation/Drug-structure

#### Data Loading and Preprocessing

The initial step involves loading multiple datasets, including RNA expression data, SMILES strings for drug compounds, drug efficacy data (IC50/AUC/resistant-sensitive values), and mutation information. These datasets are loaded into structured formats for further processing. RNA expression data is normalized and filtered to ensure consistency across samples. Missing values are handled by either imputation or removal, depending on the context. Additionally, datasets are aligned based on common identifiers, such as gene names or patient IDs, to ensure compatibility.

#### RNA Expression Data Processing

RNA expression data is processed to ensure compatibility with downstream analysis. A predefined gene list is used to filter and reindex the expression data. Genes not present in the dataset are added with zero values to maintain a consistent feature space. This step ensures that all samples have the same set of genes, which is critical for model training. The processed RNA expression data is then normalized to account for variations in sequencing depth and other technical factors.

#### Drug Fingerprint Generation

Drug compounds are represented using molecular fingerprints, which capture structural and chemical properties. SMILES strings, a textual representation of molecular structures, are converted into Morgan fingerprints using cheminformatics tools. These fingerprints are binary vectors that encode the presence or absence of specific molecular substructures. This representation allows the model to leverage structural information about the drugs, which is crucial for predicting drug efficacy.

#### Mutation Information Embedding

To incorporate mutation-specific information into the model, categorical features—such as KRAS mutation loci and metrics reflecting mutational divergence (quantified using Grantham distance)—are encoded into a fixed-dimensional vector space. An embedding matrix, initialized with random weights, maps each mutation index to a continuous representation. This transformation enables the integration of discrete genomic alterations into a differentiable model architecture. The resulting embeddings are subsequently concatenated with phenotypic and molecular features to form a unified input vector, facilitating comprehensive modeling of genotype–phenotype relationships in organoid systems.

#### Feature Concatenation

The processed RNA expression data, drug fingerprints, and mutation embeddings are combined into a single feature vector for each sample. This concatenation step integrates multiple data modalities, allowing ODFormer to leverage both molecular and genomic information. The resulting feature vectors are standardized to ensure that all features contribute equally to the model’s learning process.

#### Data Splitting and Reshaping

The combined feature vectors are split into fixed-size chunks to facilitate processing by the model. This step ensures that the input data has a consistent shape, which is necessary for efficient batch processing. The data is then divided into training and validation sets using 5-fold cross-validation. This approach ensures that the model is evaluated on multiple subsets of the data, providing a robust estimate of its performance.

### The architecture of Pretrained Dual-Encoder

#### Implementation of the Encoder Layer

To facilitate the analysis of single-cell omics data, we have designed an Encoder (Fig. 4c). The primary distinction lies in the incorporation of multiple rectified linear activation functions (ReLU) within the Transformer Decoder. Here, we adopt mask attention machine. Furthermore, the incorporation of ReLU is intended to facilitate more efficient extraction of features and enhance the learning of representations in the context of large-scale scRNA-seq data analysis.

The attention network demonstrates exceptional performance owing to several notable properties that significantly contribute to its efficacy. A salient property of the attention network is its capability to prioritize essential sub-vectors within the expression vectors. This enables ODFormer to effectively disregard the influence of insignificant sub-vectors, thereby mitigating their adverse impact on the overall analysis process. Lastly, to generate the subsequent data, a masking mechanism is employed wherein all positions after the current position are masked by setting them to infinity. This precautionary measure prevents the model from "seeing" future information. As a result, the model can solely rely on previous value to generate the next value, thereby preserving the autoregressive nature of the process. This enhances the performance of ODFormer in generating unmeasured data by ensuring that the generation process strictly adheres to the sequential dependencies and temporal coherence inherent in the data.

In addition, the employment of multiple ReLU (Rectified Linear Unit) activations confers several notable advantages. Primarily, ReLU is computationally efficient. Unlike more complex activation functions such as sigmoid or hyperbolic tangent (tanh), ReLU performs a straightforward thresholding operation wherein negative inputs are mapped to zero. This simplicity in computation accelerates the pretraining process of ODFormer, enhancing their ability to generate outputs that closely approximate measured data. Another significant benefit of ReLU is its capacity to introduce sparsity into activations. This sparsity facilitates more efficient information processing by encouraging ODFormer to concentrate on pertinent features while disregarding irrelevant or noisy inputs, which may arise from various factors including sequencing technicalities, low-confidence reads, batch effects, and cell capture biases. Additionally, ReLU contributes to mitigating the overfitting problem by reducing the model’s tendency to excessively focus on noise or outliers in the data. This inherent regularization effect enables ODFormer to achieve exceptional performance even with limited annotated scRNA-seq data.

Attention network in ODFormer is sc-attention network, calculated by the following operation:

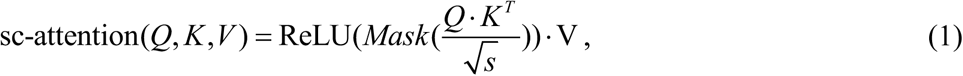

where *Q=*ReLU*(W^q^X)*, *K=*ReLU*(W^k^X*_posi_), *V=*ReLU*(W^v^X*_posi_), the *W*^q,k,v^ is the linear project weight, ReLU(*z_i_*) = max(*z_i_*, 0).

For ODFormer, we use multi-head self-attention to capture different interaction information in multiple different projection spaces. We set the hyper parameter head to 32 in ODFormer, which means we have 32 parallel self-attention operations, i.e.

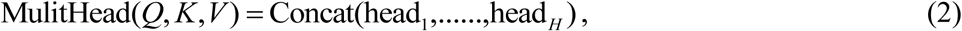

where

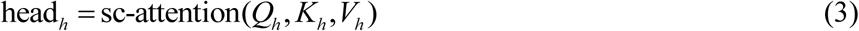

After layer normalization and multi-head attention,

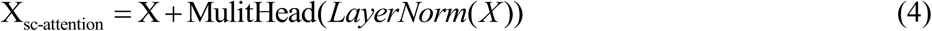

where *X* is converted into *X_sc-_*_attention_ as the input of the feed-forward network.

The sc-attention network is useful for capturing dependencies between elements among sub-vectors. However, it may not be enough to capture all non-linear features among sub-vectors. To address this, the feed-forward network is added. Feed-forward network consists of two linear layers, with a ReLU between them, i.e.

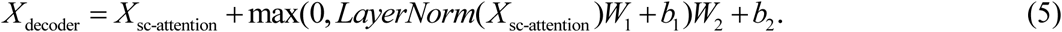

#### Implementation of the Deconder Layer

Preceding the Linear Layer, the application of an average pooling layer is employed to compute the mean value across all sub-vectors.

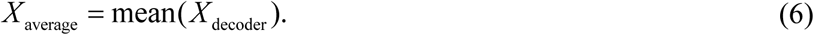

We then use one linearly connected network to reconstruct the cell

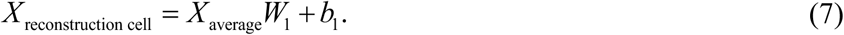

#### Loss Function

To achieve a closer resemblance of the reconstructed data to the measured data, we employ a MSE loss function. Within this function, *X* represents the measured cell data, while *X*_reconstruction_ represents the output generated by encoder:

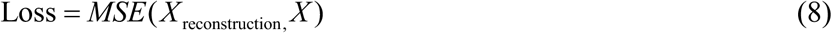

### The architecture of downstream task model of ODFormer

#### Model Architecture Overview

The Transformer-based model described here is designed to handle sequential or structured data, making it well-suited for drug response prediction tasks. The architecture leverages the Transformer encoder, which consists of multiple layers of self-attention and feed-forward neural networks. These components enable the model to capture complex dependencies within the input data, such as interactions between drug properties and biological context. The model begins by processing the input data, which is typically represented as a multi-dimensional array with dimensions corresponding to batch size, sequence length, and feature dimensions. Positional encoding is added to the input data to provide information about the order of elements in the sequence, addressing the Transformer’s lack of inherent sequential awareness. The encoded data is then passed through the Transformer encoder, which generates enriched representations of the input. Finally, the model applies a pooling operation to produce a fixed-size representation of the sequence, which is used to make predictions. This flexible architecture allows the model to be adapted for both regression and classification tasks, depending on the specific requirements of the drug response prediction problem.

#### Regression Model for Continuous Drug Response Metrics

When the goal is to predict continuous drug response metrics such as IC50 (half-maximal inhibitory concentration) or AUC (area under the dose-response curve), the model is configured as a regression model. These metrics quantify the potency or efficacy of a drug against a specific biological target or cell line. The input data consists of features representing drug properties, such as molecular descriptors or chemical fingerprints, and biological context, such as gene expression profiles or mutation status. These features are structured as sequences or fixed-length vectors. The model processes the input data through the Transformer encoder, which captures complex relationships between the features. The output of the encoder is pooled to produce a fixed-size representation of the sequence, which is then passed through a series of fully connected layers to generate the final prediction. The output layer uses a linear activation function to produce unbounded predictions, and the model is trained using mean squared error (MSE) or mean absolute error (MAE) as the loss function. This configuration allows the model to accurately predict continuous drug response metrics, providing valuable insights into drug potency and efficacy.

#### Binary Classification Model for Resistance/Sensitivity Prediction

When the goal is to predict categorical outcomes such as resistant or sensitive, the model is configured as a binary classification model. This approach is useful for determining whether a drug is likely to be effective against a specific cell line or patient sample. The input data is similar to the regression model, consisting of features representing drug properties and biological context, but the target variable is binary, with 0 indicating resistance and 1 indicating sensitivity. The model processes the input data through the Transformer encoder, which captures complex dependencies between the features. The output of the encoder is pooled to produce a fixed-size representation of the sequence, which is then passed through a series of fully connected layers to generate the final prediction. The output layer uses a sigmoid activation function to produce a probability score between 0 and 1, indicating the likelihood of sensitivity. The model is trained using binary cross-entropy loss, which is well-suited for binary classification tasks. This configuration enables the model to classify drug-cell line pairs into resistant or sensitive categories, providing actionable insights for drug selection and patient stratification.

#### Hybrid Contrastive Loss

The HybridContrastiveLoss function is an innovative loss function that aims to optimize multi-modal representations by combining two crucial contrastive learning strategies: the Adaptive Contrastive Loss and the Distance Contrastive Loss. These two components help the model learn meaningful embeddings that capture both the inherent structure of patient data and the relationships between patients, drugs, and treatment responses. The loss function’s unique aspect lies in its adaptive mechanisms, which adjust the contribution of each component based on training dynamics.

The Adaptive Contrastive Loss utilizes a triplet loss mechanism, which is designed to learn embeddings that bring similar samples (anchor and positive) closer together while pushing dissimilar samples (anchor and negative) farther apart. This approach is particularly effective for embedding tasks where the goal is to learn a feature space that maintains meaningful relationships between the samples. A triplet consists of an anchor, a positive sample (similar to the anchor), and a negative sample (dissimilar to the anchor). The triplet loss can be expressed mathematically as:

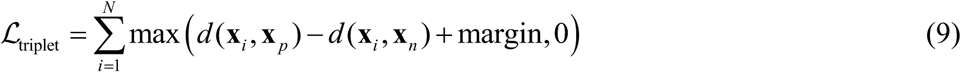

Here, *x_i_* is the anchor embedding (e.g., a patient’s embedding), *x_p_* is the positive sample embedding (a patient that is similar to the anchor), *x_n_* is the negative sample embedding (a patient that is dissimilar to the anchor), *d* ( *x_i_*, *x_p_*) and *d* ( *x_i_*, *x_p_*) represent the distances between the anchor and the positive sample, and between the anchor and the negative sample, respectively, The margin is a fixed threshold that enforces a minimum separation between positive and negative pairs.

The loss ensures that the model learns a feature space where the anchor and positive pair are closer in distance than the anchor and negative pair by at least the margin value. To address the issue of class imbalance in triplet selection, the Adaptive Contrastive Loss introduces dynamic thresholding for response and drug similarities. Specifically, the thresholds for these similarities are adjusted based on the number of positive and negative samples in a given batch. This adaptive thresholding allows the model to continuously adjust its sample selection strategy throughout the training process, ensuring that the learning process remains balanced and effective.

The adaptive thresholds for response and drug similarity are updated as follows:

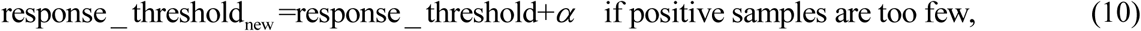

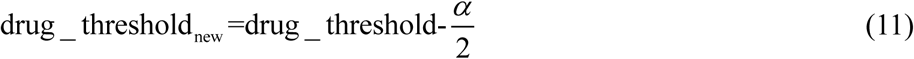

where α is a learning factor that determines how much the threshold should be adjusted. Conversely, when negative samples are too few, the thresholds are adjusted in the opposite direction:

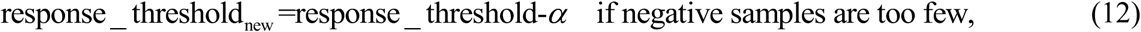

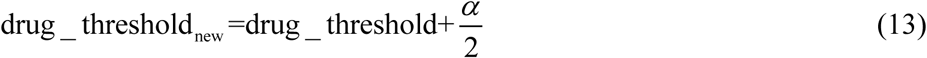

The Distance Contrastive Loss aims to align the learned patient embeddings with the external relationships between drugs and patient responses. This is particularly useful for tasks such as predicting drug efficacy or response rates, where the relationships between patient features, drug properties, and response behaviors are crucial.

The Distance Contrastive Loss is calculated by measuring the cosine similarity between patient embeddings and comparing this with the target similarity, which is derived from the drug and response embeddings. The cosine similarity between two vectors aa and bb is given by:

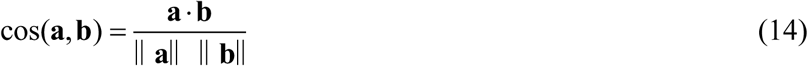

For the patient embeddings *p_i_* and *p_j_*, the cosine similarity is:

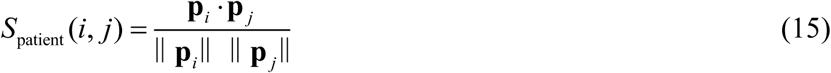

The target similarity *S*_target_ (*i*, *j*) is derived from the combination of the drug and response similarities, which are modeled separately. The drug similarity is based on cosine similarity between drug embeddings, and the response similarity is computed based on the difference between response values. The target similarity is the product of these two similarities:

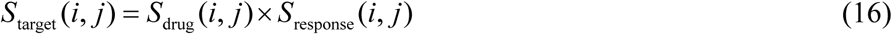

The Distance Contrastive Loss is then computed as the mean squared error (MSE) between the patient similarity *S*_patient_ (*i*, *j*) and the target similarity *S*_target_ (*i*, *j*) :

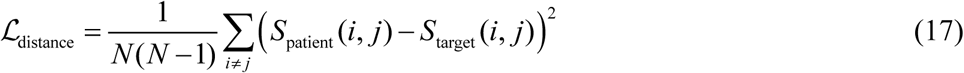

The HybridContrastiveLoss function combines both the Adaptive Contrastive Loss and the Distance Contrastive Loss in a way that dynamically adjusts their contributions during training. This hybrid approach enables the model to learn robust, high-quality representations by optimizing for both internal consistency (through the triplet loss) and external alignment (through the distance contrastive loss).

The final HybridContrastiveLoss is computed as a weighted sum of the two losses:

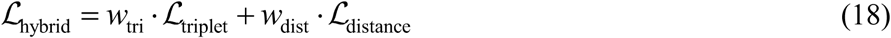

Where *w*_tri_ and *w*_dist_ are adaptive weights that are computed dynamically based on the relative magnitudes of the individual losses. These weights ensure that the model balances the contributions of the two loss components according to the training dynamics. If one loss component dominates, the model may shift focus toward the other component accordingly.

The weight adjustment is based on the ratio of the losses, such that:

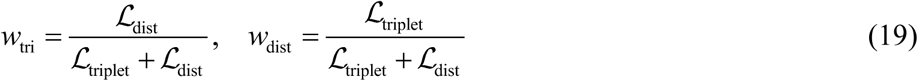

This dynamic adjustment helps the model focus on the most important aspects of the learning process at any given stage, ensuring that both triplet-based and distance-based losses contribute effectively to the optimization.

The HybridContrastiveLoss function is highly effective for multi-modal learning tasks, especially those where there are complex interdependencies between features, such as in medical data analysis. By combining the strengths of both triplet loss and distance contrastive loss, this hybrid approach ensures that the model learns meaningful, high-quality embeddings that capture both intra-modal and inter-modal relationships.

The adaptive thresholding mechanism ensures that the model is not overly biased towards either positive or negative samples, preventing the model from overfitting or underfitting. This dynamic adjustment allows the model to handle imbalanced data distributions and adapt to changes in the training process.

Furthermore, the HybridContrastiveLoss ensures that the learned embeddings are both internally consistent and externally aligned with the relationships between drugs, responses, and patient characteristics, making it particularly well-suited for tasks like drug response prediction and precision medicine.

In summary, the HybridContrastiveLoss provides a flexible, adaptive framework that optimizes multi-modal embeddings for complex, real-world tasks. By integrating both triplet-based and distance contrastive losses, it offers a powerful tool for learning rich, informative representations in settings where multiple feature types must be considered simultaneously.

### Evaluation metrics

To systematically assess the model’s predictive performance, we employed a comprehensive set of metrics appropriate to each task. For regression tasks, we quantified prediction accuracy using the Pearson correlation coefficient, which evaluates the linear relationship between predicted values *x*_i_ and ground-truth values *y*_i_. The PCC is defined as:

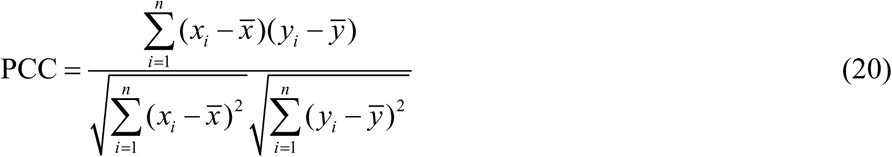

For ranking-based evaluations, we adopted four standard metrics: mean average precision (MAP), normalized discounted cumulative gain (NDCG), mean reciprocal rank (MRR), and hit rate@K. MAP calculates the mean of average precision across queries:

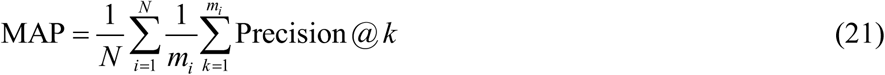

NDCG at rank *K* is given by:

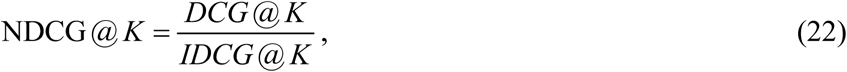

where

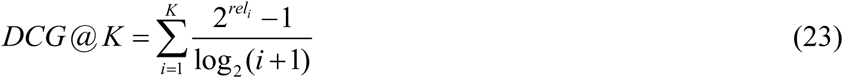

MRR measures the average reciprocal rank of the first relevant item:

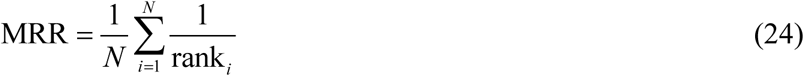

Hit rate evaluates the proportion of samples where a relevant item is found within the top-KKK predictions:

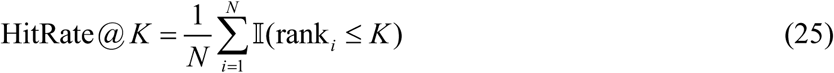

For classification tasks, we used three complementary metrics: accuracy (ACC), area under the receiver operating characteristic curve (ROAUC), and area under the precision-recall curve (PRAUC). Accuracy is calculated as:

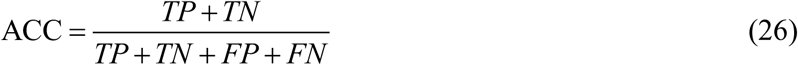

ROAUC and PRAUC are computed as the area under their respective performance curves:

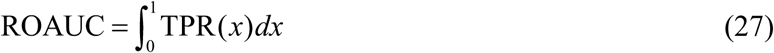

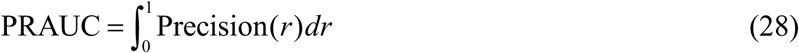

These metrics provide a robust assessment of model performance across diverse prediction scenarios.

### Data Collection and pre-processing

#### Samples and patients

Tumor samples were collected from pancreatic cancer patients who received surgical resection or EUS-FNA at Changhai Hospital. Samples were pathologically confirmed as tumor or normal tissue. Clinical variables of patients were collected from the prospective database of Changhai Hospital. All patients provided written informed consent for the use of their clinical data and surgical specimens, and consent to publish clinical information potentially identifying individuals was obtained. The study was conducted in accordance with the national guidelines and was approved by the Ethics Committee of Changhai Hospital (Approval number: CHEC2021-055).

#### Organoid culture

The following culture media were used: basic medium (advanced DMEM/F12, 10 mM HEPES, 1X GlutaMAX-I, 100 µg/ml Primocin, 1X penicillin/streptomycin solution) and complete medium (advanced DMEM/F12, 10 mM HEPES, 1X GlutaMAX-I, 100 µg/ml Primocin, 1X penicillin/streptomycin solution, 500 nM A83-01, 10 µM Y-27632, 1.56 mM N-acetylcysteine, 10 mM nicotinamide, 10 ng/ml FGF10, 1X B27 supplement, 10 µM forskolin, 30% Wnt3A conditioned medium, 2% R-spondin conditioned medium, 4% Noggin conditioned medium). For surgical samples, tissues were minced and incubated in digestion medium (2.5 mg/ml collagenase II and 10 µM Y-27632 in basic medium) at 37 °C with mild agitation for approximately 1 h. The obtained cells were cultured on suspension plates with Matrigel and complete medium (PDAC samples were cultured in complete medium, while samples of normal pancreas and other subtypes of pancreatic cancer were cultured in complete medium supplemented with EGF 50 ng/ml). Material obtained from biopsy samples was directly cultured under the above conditions without digestion. The media used for organoid cryopreservation were composed of the corresponding culture medium (90%) and 10% DMSO. The established organoids were routinely tested for mycoplasma contamination. All organoid experiments were performed at the Shanghai Institute of Biochemistry and Cell Biology^29^.

#### Whole-genome library preparation and sequencing

Genomic DNA from PDPCOs and blood was extracted using a QIAamp DNA kit (Qiagen, 51306). Whole-genome libraries were generated using a NEBNext Ultra DNA Library Prep Kit (New England Biolabs, E7370L) according to the manufacturer’s protocols. A 1 μg aliquot of DNA was used as input for fragmentation into ∼300 bp fragments using a Covaris LE220 sonicator, and purification was performed with DNA Clean Beads. The DNA fragments underwent bead-based size selection and subsequent end repair, adenylation, and ligation to Illumina sequencing adapters. The final libraries were evaluated using a QIAxcel bioanalyzer. Libraries were sequenced on the Illumina HiSeq XTen platform aiming for 30X and 50X coverage for blood and organoids, respectively, and 150 bp paired-end reads were generated.

#### Organoids RNA library preparation and sequencing

PDPCOs and NPOs in Matrigel were collected and washed with precooled PBS before lysis with 1 ml of TRIzol (Invitrogen, 15596018), and total RNA was extracted according to the manufacturer’s instructions. A total amount of 3 μg RNA per sample was used as input material for RNA sample preparation. Ribosomal RNA was removed with a Globin-Zero Gold rRNA Removal Kit (Illumina, E7750X), and rRNA-free residue was cleaned up by ethanol precipitation. RNA-seq libraries were generated using a NEBNext Ultra RNA Library Prep Kit (New England Biolabs, E7530L) according to the manufacturer’s instructions. After cluster generation, the libraries were sequenced on the Illumina HiSeq X Ten platform, and 150 bp paired-end reads were generated^29^.

#### Chemotherapy Regimen Recommendation Based on Organoid/ODFormer Sensitivity Prediction

To optimize the selection of chemotherapy agents, we developed a percentile-based stratification approach for drug sensitivity predictions, integrating in vitro organoid testing with ODFormer computational modeling. Sensitivity scores for each chemotherapeutic agent were derived from patient-specific data and ranked in ascending order. Based on these rankings, agents were categorized into three distinct tiers: sensitive (sen), intermediate (mid), and resistant (res). The top 30% of agents, which showed the highest predicted efficacy, were classified as sensitive, while the bottom 30% were categorized as resistant, indicating the lowest predicted efficacy. The remaining 40% were classified as intermediate, reflecting moderate predicted efficacy.

The chemotherapy regimen for each patient was then prioritized according to these classifications. Agents classified as sensitive were recommended as the primary treatment option, ensuring the highest therapeutic potential. In cases where no sensitive agents were available, those in the intermediate tier were considered as secondary options. Resistant agents, with their low predicted efficacy, were systematically excluded from the regimen to avoid the risk of administering ineffective cytotoxic treatments.

This tiered recommendation system provides a personalized approach to chemotherapy, enhancing the likelihood of successful treatment while minimizing the exposure to ineffective agents, ultimately improving patient outcomes.

#### Tissues RNA library preparation and sequencing

Total RNA was isolated from tumor tissues using the TRIzol reagent (Invitrogen, Carlsbad, CA, USA) and purified using the RNAeasy Mini Kit (Qiagen, Hilden, Germany) followed by the manufacturer’s instructions. RNA degradation and contamination was assessed via agarose gel electrophoresis. RNA purity was measured using NanoPhotometer® spectrophotometer (IMPLEN, CA, USA) and RNA integrity was confirmed by the RNA Nano 6000 Assay Kit of the Bioanalyzer 2100 system (Agilent Technologies, CA, USA). At least 1 μg of total RNA per sample was used as input material for RNA library preparation. Stranded RNA-seq libraries were constructed using the NEB- Next® UltraTM RNA Library Prep Kit for Illumina® (NEB, USA) following the manufacturer’s instructions. Briefly, mRNA was enriched from total RNA by oligo dT magnetic beads and fragmentation. Then, synthesis of first-strand cDNA was performed by using random hexamer primer and followed by synthesis of second-strand cDNA using DNA Polymerase I and RNase H. Fragments size selection, adaptor ligation as well as PCR amplification were performed subsequently. The cDNA library quality was evaluated on the Agilent Bioanalyzer 2100 system. The index-coded samples were carried out on a cBot Cluster Generation System using Illumina PE Cluster Kit (Illumina, San Diego, California, USA) for cluster generation, and cDNA libraries were sequenced on an Illumina Hiseq XTEN platform to produce 150bp paired-end reads (Illumina, San Diego, California, USA)

#### High-throughput screening of chemical and chemotherapeutic drugs

White, clear bottom 384-well plates were coated with 10 μl of collagen at room temperature for at least one hour using a Multidrop Combi reagent dispenser before the addition of organoid suspensions. PDPCOs were dissociated with Tryp-LE before being resuspended in medium and dispensed into 384-well plates (3,500 cells per well). The next day, 283 compounds (Selleck), as well as DMSO controls were added in duplicate using a Bravo robotic workstation. To assess cell viability, 25 μl of CellTiter-Glo Reagent per well was added after three days. The plates were gently shaken for 15 min at room temperature before luminescence was assessed using an Envision plate reader. Average inhibition rates from two independent experiments were calculated with Excel and visualized using GraphPad Prism 8.

Five chemotherapeutic agents (gemcitabine, paclitaxel, 5-fluorouracil, oxaliplatin and irinotecan) and chemicals that had significant inhibitory effects on cells were used for the secondary screening. The range of concentrations selected for each chemical was based on the primary screening data. Organoids were similarly dispensed into 384-well plates. Concentration dilution and addition of each compound were performed with the Bravo robotic workstation. Cell viability was assessed using CellTiter-Glo Reagent after three days of incubation with drugs. The secondary screening was performed in technical duplicate (same screening run), and all screening plates were subjected to stringent quality control measures. To measure sensitivity, we used 5-point dose-response curves; for each drug, was 5 concentrations and the corresponding cell viability values were used as input for curve generation. The viability was set to 100 if it was higher than baseline, and each drug concentration (nmol/L) was log10 transformed. The AUC was calculated with the sintegral function in R, and the normalized AUC was obtained by dividing one AUC by the maximum AUC for each drug. The AUC heatmap for the secondary screening was generated with GraphPad Prism 8.

#### Clinical follow-up

Clinical follow-up data were collected from the Changhai Hospital prospective database. Patients regularly received tumor marker and imaging examinations every three months after surgery and were followed up at the same time interval by outpatient clinic visits and/or telephone contact.

#### Survival analysis

Survival analysis was performed to assess the time to event (or failure) for each subject in the dataset. The analysis utilized the Kaplan-Meier estimator for estimating the survival function and the Cox Proportional-Hazards model to examine the influence of covariates on survival time.

The Kaplan-Meier estimator was applied to estimate the survival probability over time. The survival function S(t) was calculated for each individual, and the event data was represented as 1 for the occurrence of the event and 0 for censored data. Censoring occurred when a subject did not experience the event during the observation period. The Kaplan-Meier estimator was computed using the lifelines library in Python, and the survival curve was plotted using the KaplanMeierFitter function.

#### Transcriptome Data Preprocessing

This refers to the steps involved in preparing RNA sequencing data for analysis, such as quality control, filtering out low-quality reads, removing contaminants, and normalizing expression levels.

#### Single-Cell Transcriptome Data Preprocessing

This involves preparing single-cell RNA sequencing data by handling issues like cell quality filtering, removing unwanted noise, correcting for batch effects, and normalizing data for accurate down-stream analysis.

#### AUC Standardized Data Preprocessing

This process involves preparing drug sensitivity data, AUC (area under the curve) values, by normalizing and transforming the data to ensure consistency and accuracy in comparative analyses across experiments.

### Tools for comparison

**DeepCDR** combines multi-omics data (such as genomic and transcriptomic profiles) of cancer cells with the chemical structure information of drugs using a hybrid graph convolutional network (GCN). This method incorporates both a uniform GCN and multiple subnetworks to automatically learn the latent representations of drug structures, unlike traditional methods that rely on handcrafted drug features. By doing so, DeepCDR provides more accurate predictions of cancer drug responses, demonstrating superior performance in both classification and regression tasks.

**BANDRP** utilizes a bilinear attention model that integrates gene expression data of cancer cell lines with molecular fingerprints of drugs to predict cancer drug responses. It calculates pathway enrichment scores to enhance the features of cancer cell lines and learns the interactions between cancer cell lines and drugs through bilinear attention networks. BANDRP efficiently captures the interactive information between cancer cell lines and drugs, outperforming baseline models in drug response prediction.

**PANCDR** is an adversarial network-based model designed to bridge the gap between preclinical and clinical datasets. The method includes an adversarial model to reduce dataset discrepancies and a cancer drug response (CDR) prediction model that estimates drug responses based on preclinical data and unlabeled clinical data. PANCDR offers an important tool for precision medicine, providing personalized drug recommendations and outperforming other machine learning models in predicting cancer drug responses.

## Data Availability

All datasets used in this study are publicly available and the usages are fully illustrated in the Supplementary Table.

## Code Availability

ODFormer is implemented using PyTorch with code available at https://github.com/xujing363/ODFormer.

## Author contributions

L.C., G.J., D.G., J.X. and X.Y. conceived and designed the project. J.X. developed and implemented the computational framework, and performed benchmarks and case studies under the guidance of L.C. and G.J. Clinical data collection and organization were carried out by X.Y. Data analysis was performed by J.X. and X.Y. Organoid drug sensitivity experiments were conducted by Y.L., H.W., and Y.L. Drug sensitivity data curation was done by S.T. Result interpretation was performed by J.X and X.Y. The manuscript was written by J.X. and X.Y., with contributions from L.C., G.J., and D.G. All authors read and approved the final manuscript.

## Acknowledgements

This work has been supported by National Key R&D Program of China (2022YFA1004800, 2025YFF1207900), Natural Science Foundation of China (T2341007, T2350003, 12131020, 42450084, 42450135, 12326614, and 12426310), Science and Technology Commission of Shanghai Municipality (23JS1401300), Zhejiang Province Vanguard Goose-Leading Initiative (2025C01114), Hangzhou Institute for advanced study of UCAS (2024HIAS-P004), and JST Moonshot R&D (JPMJMS2021).

