## Supplementary for "ODFormer: a Virtual Organoid for Predicting Personalized Therapeutic Responses in Pancreatic Cancer"

### Supplementary Figures

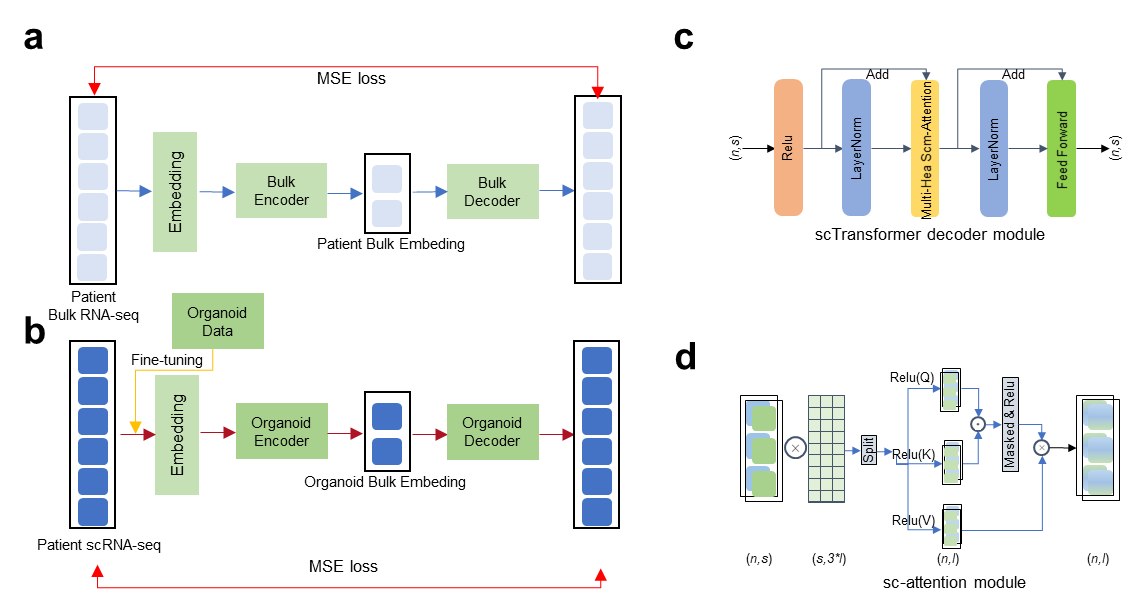

#### Supplementary Figure 1: Model of Pretrained Model

This illustrates a dual-modality sequence-to-sequence deep learning framework designed to model and reconstruct transcriptional patterns from both bulk and organoid RNA-seq data. The architecture consists of two parallel autoencoder pipelines—one for bulk data and the other for organoid data. In each pipeline, the input sequences are first transformed into high-dimensional embeddings, which are then processed by modality-specific Transformer encoders and decoders to capture contextual dependencies and latent structures.

In the bulk branch, the input undergoes embedding, followed by encoding through a Bulk Encoder. The resulting latent representation is subsequently decoded by the Bulk Decoder to reconstruct the original input. Similarly, the organoid branch processes its input through an Embedding module and an Organoid Encoder, and integrates additional organoid-specific metadata before passing it through the Organoid Decoder for reconstruction.

The top-right inset provides a detailed view of the internal structure of the Transformer blocks used in both encoders and decoders, highlighting key components such as Multi-Head Self-Attention, Layer Normalization, ReLU activation, and Feed Forward layers. The bottom-right inset further illustrates the mechanics of the self-attention operation, including query-key dot product computation, attention score normalization, and masked ReLU activation.

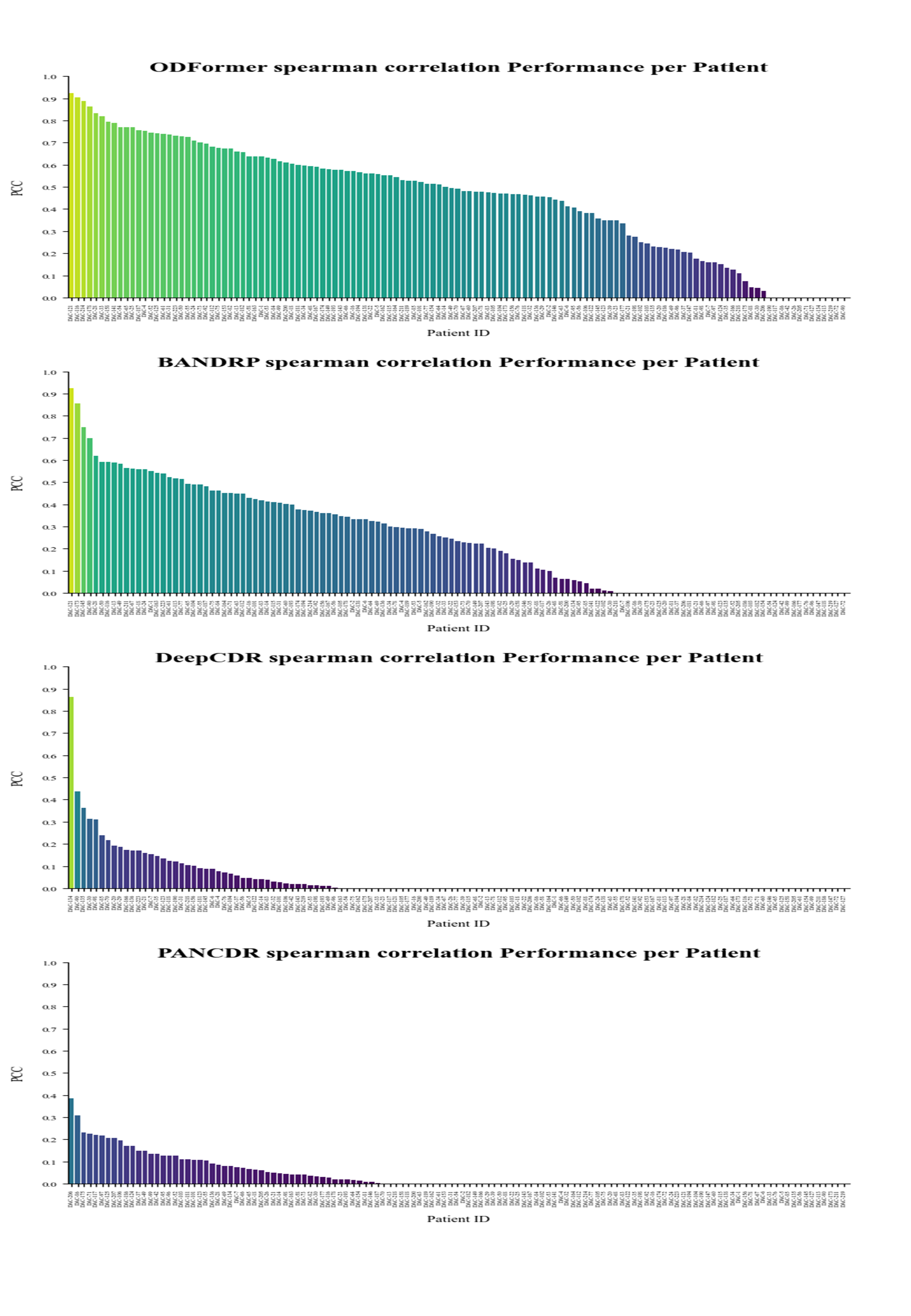

#### Supplementary Figure 2: Performance of methods in individual patient.

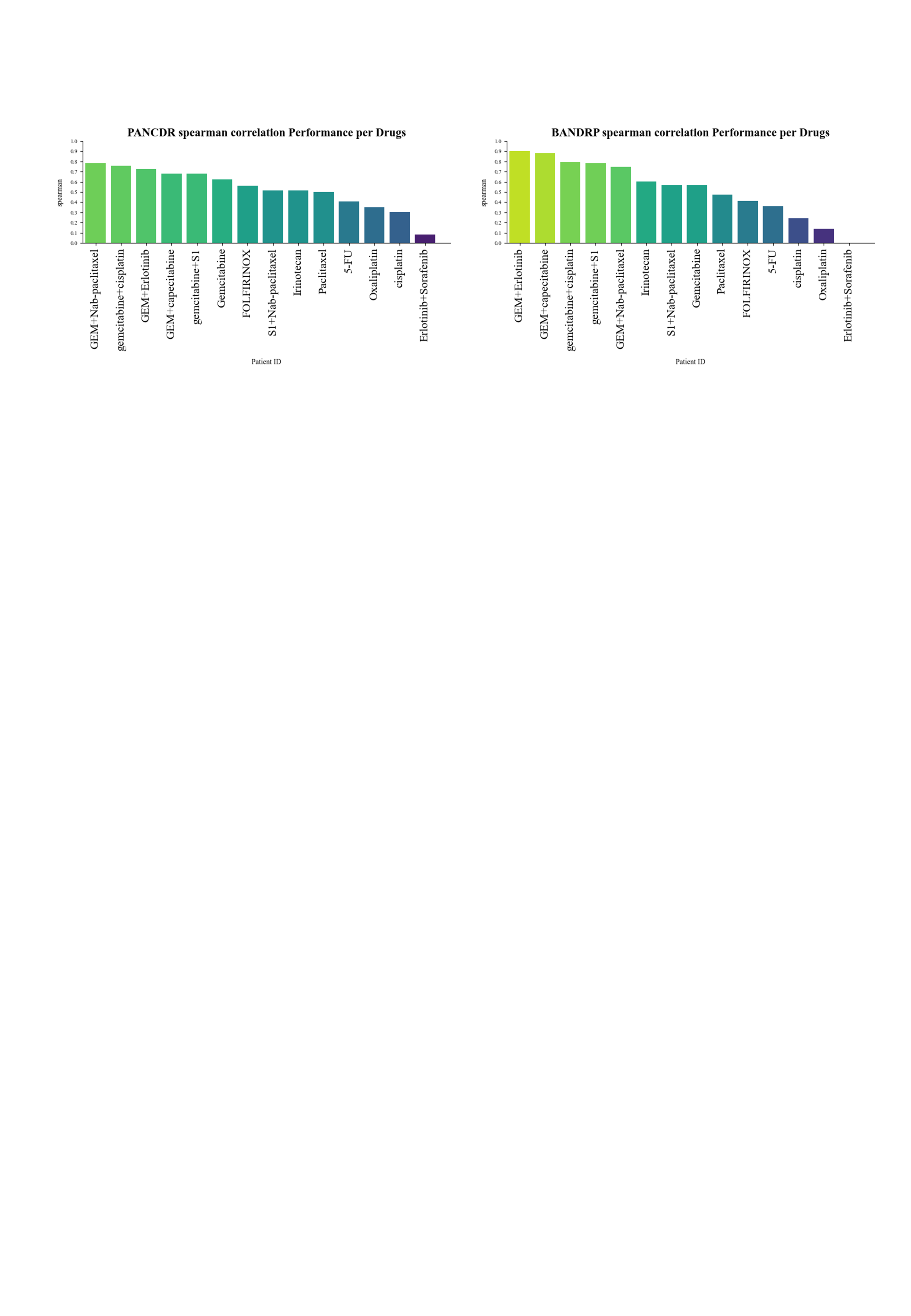

#### Supplementary Figure 3: Performance of methods in individual drugs..

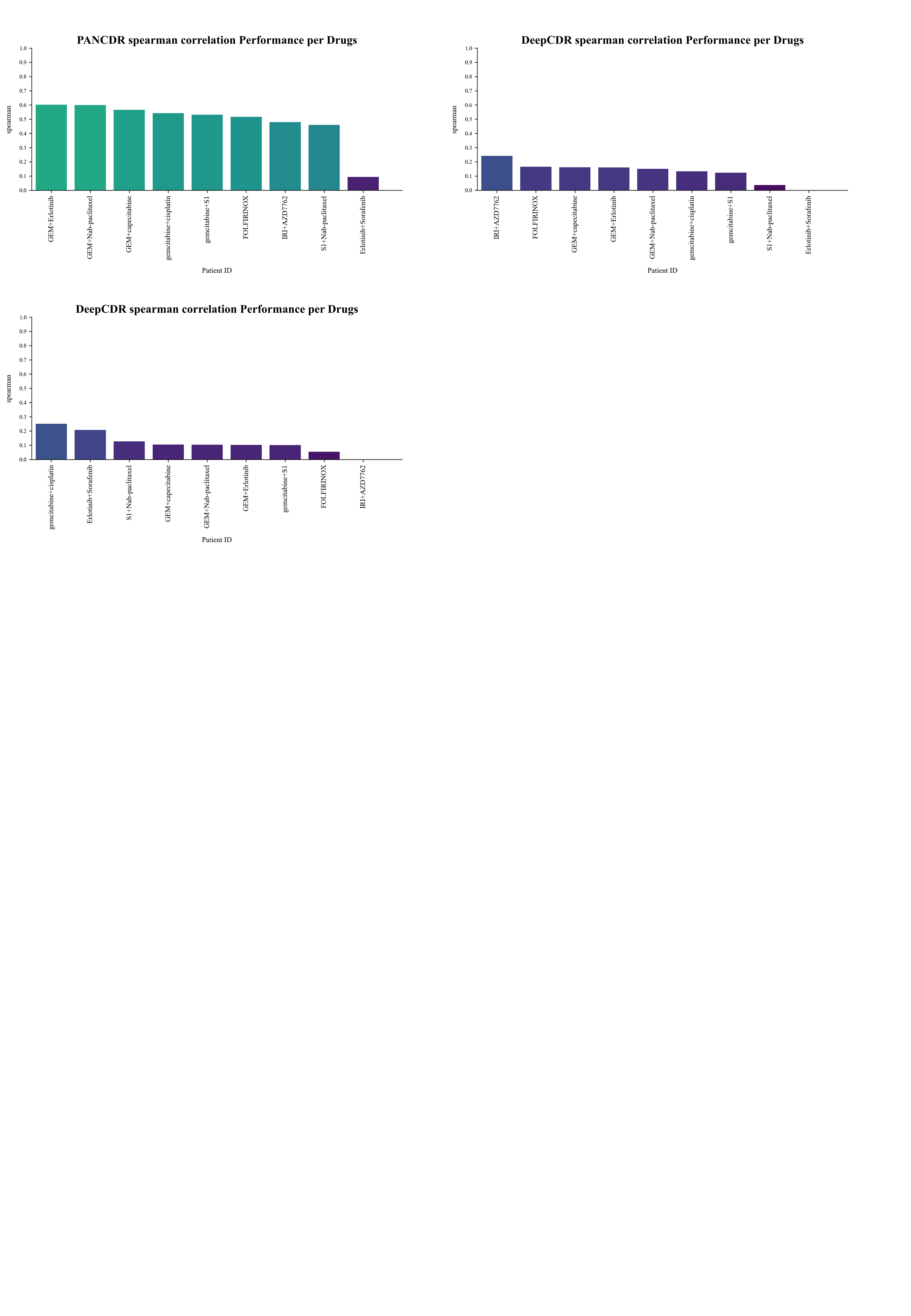

#### Supplementary Figure 5: Performance of methods in individual drugs in the task of **predicting responses to** novel drug combinations **using only monotherapy data**.

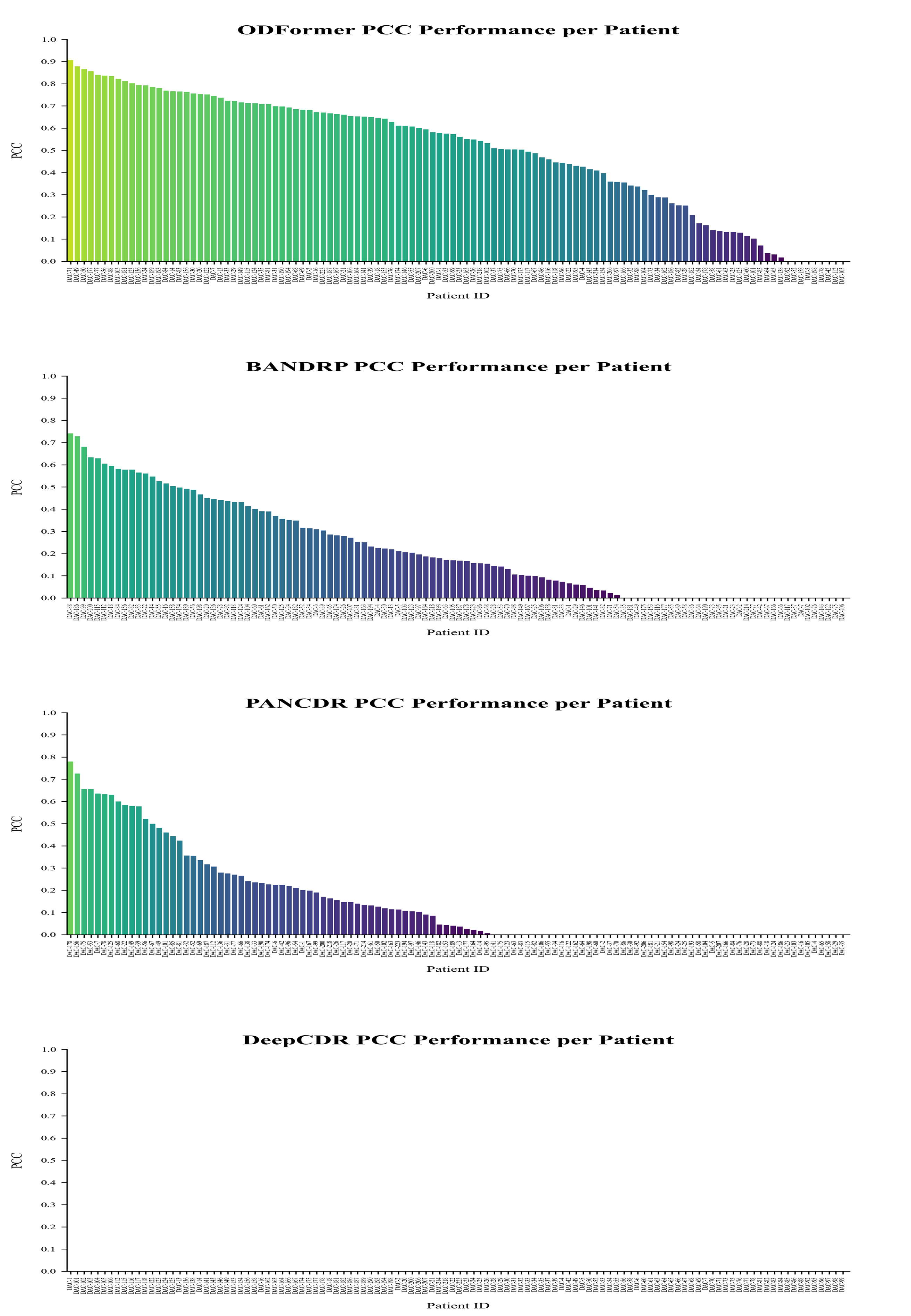

#### Supplementary Figure 6: Performance of methods in individual patients in the task of **predicting responses to** novel drug combinations **using only monotherapy data**.

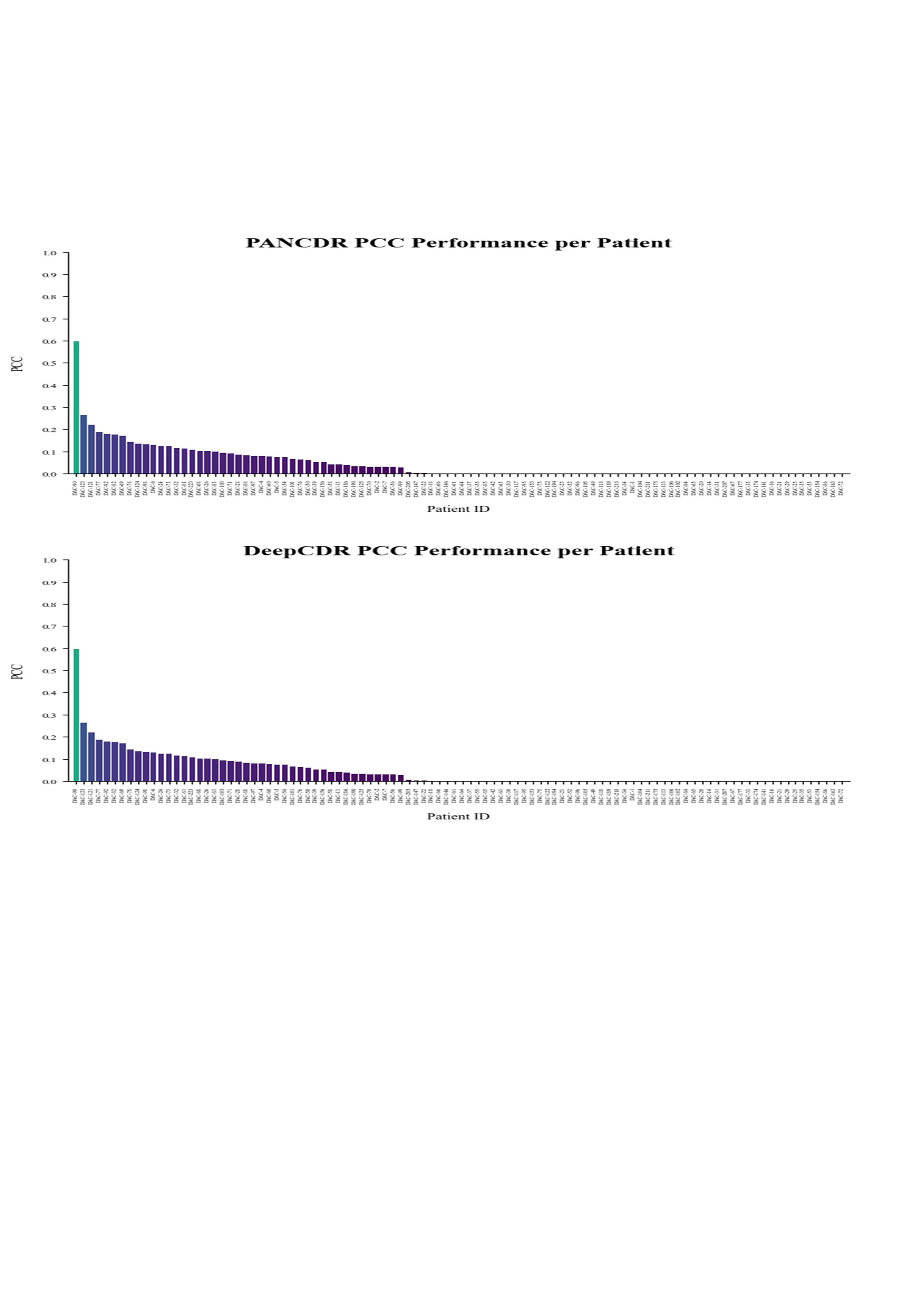

#### Supplementary Figure 7: Performance of methods in individual patients in the task of simulating the generation of virtual organoids and predicted DR-AUC directly from patient tumor tissue transcriptomes.

.

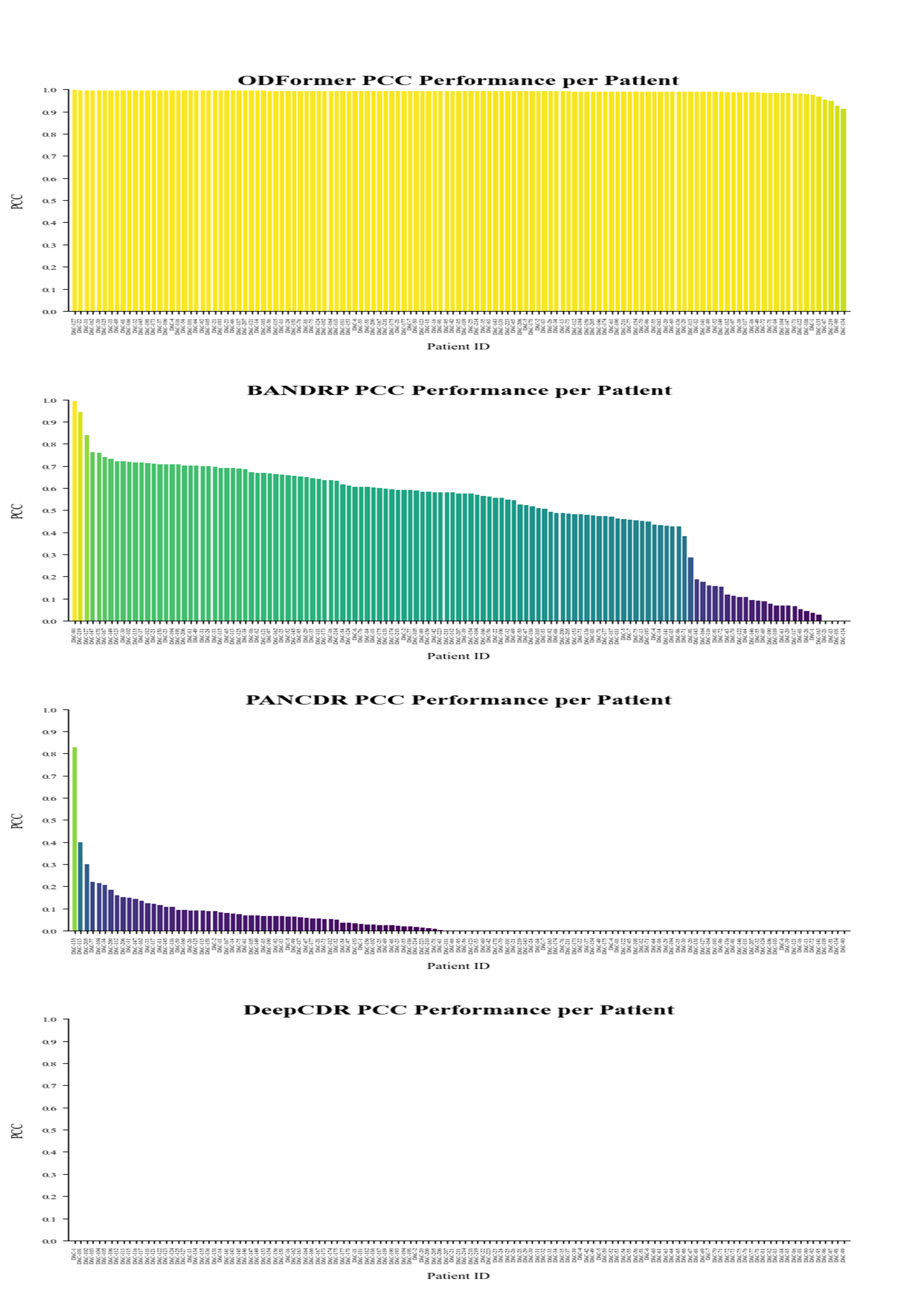

#### Supplementary Figure 8: Performance of methods in individual patients in the task of simulating the generation of virtual organoids and predicted Normalized DR-AUC directly from patient tumor tissue transcriptomes.

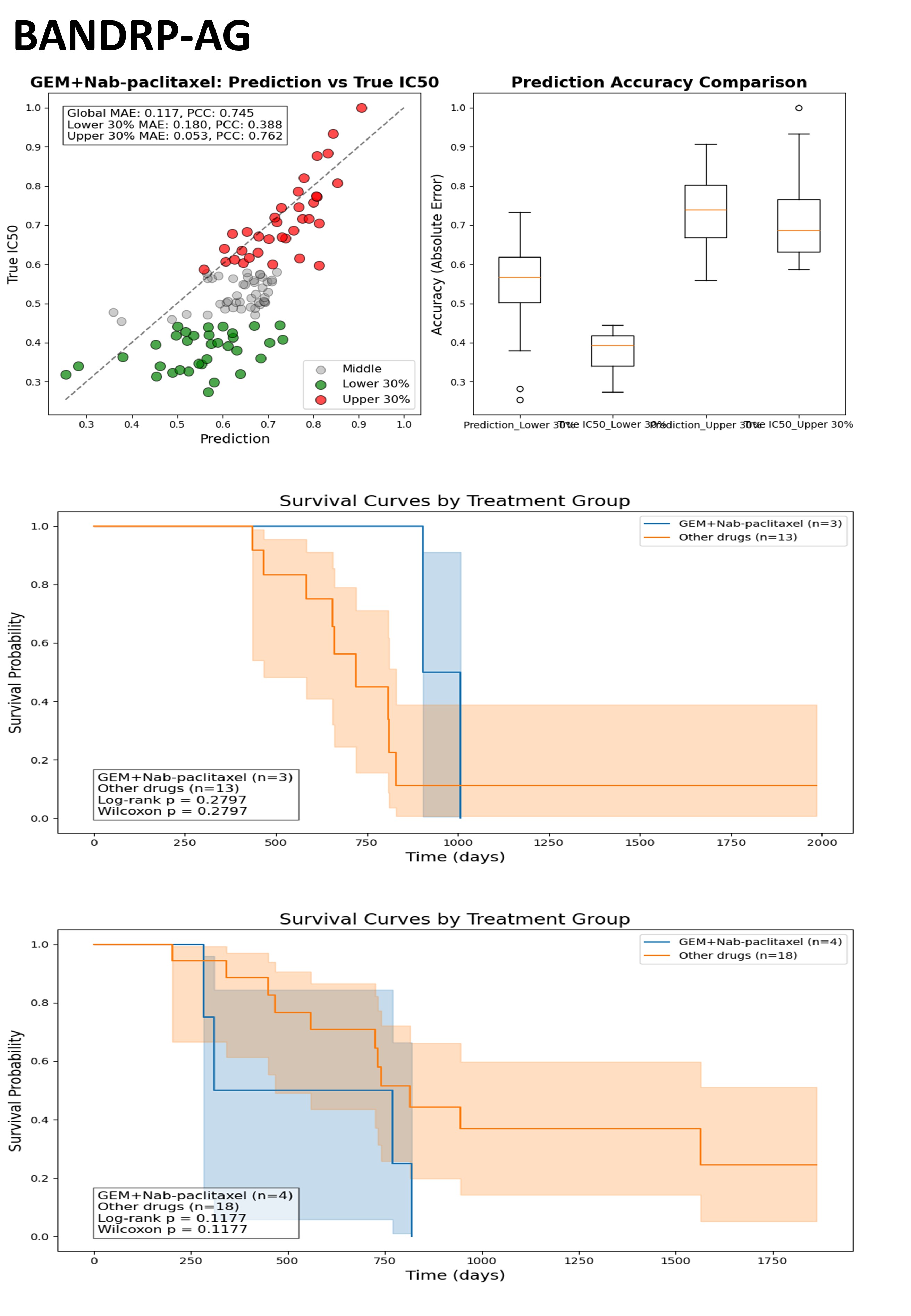

#### Supplementary Figrue 9: Performance of BANDRP-guided AG Personalized Treatment Recommendations Im-proves Patient Overall Survival.

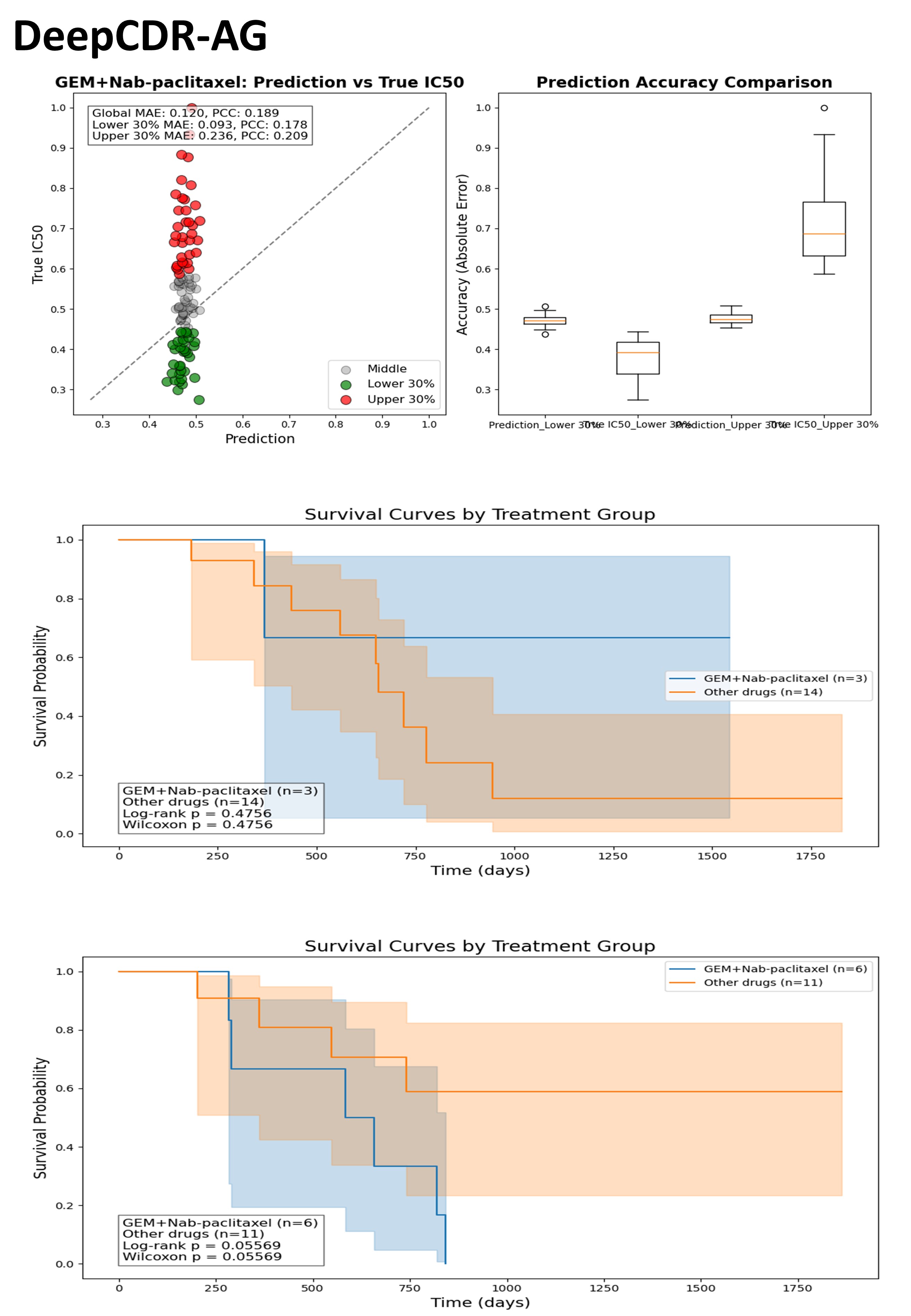

#### Supplementary Figure 10: Performance of DeepCDR-guided AG Personalized Treatment Recommendations Im-proves Patient Overall Survival.

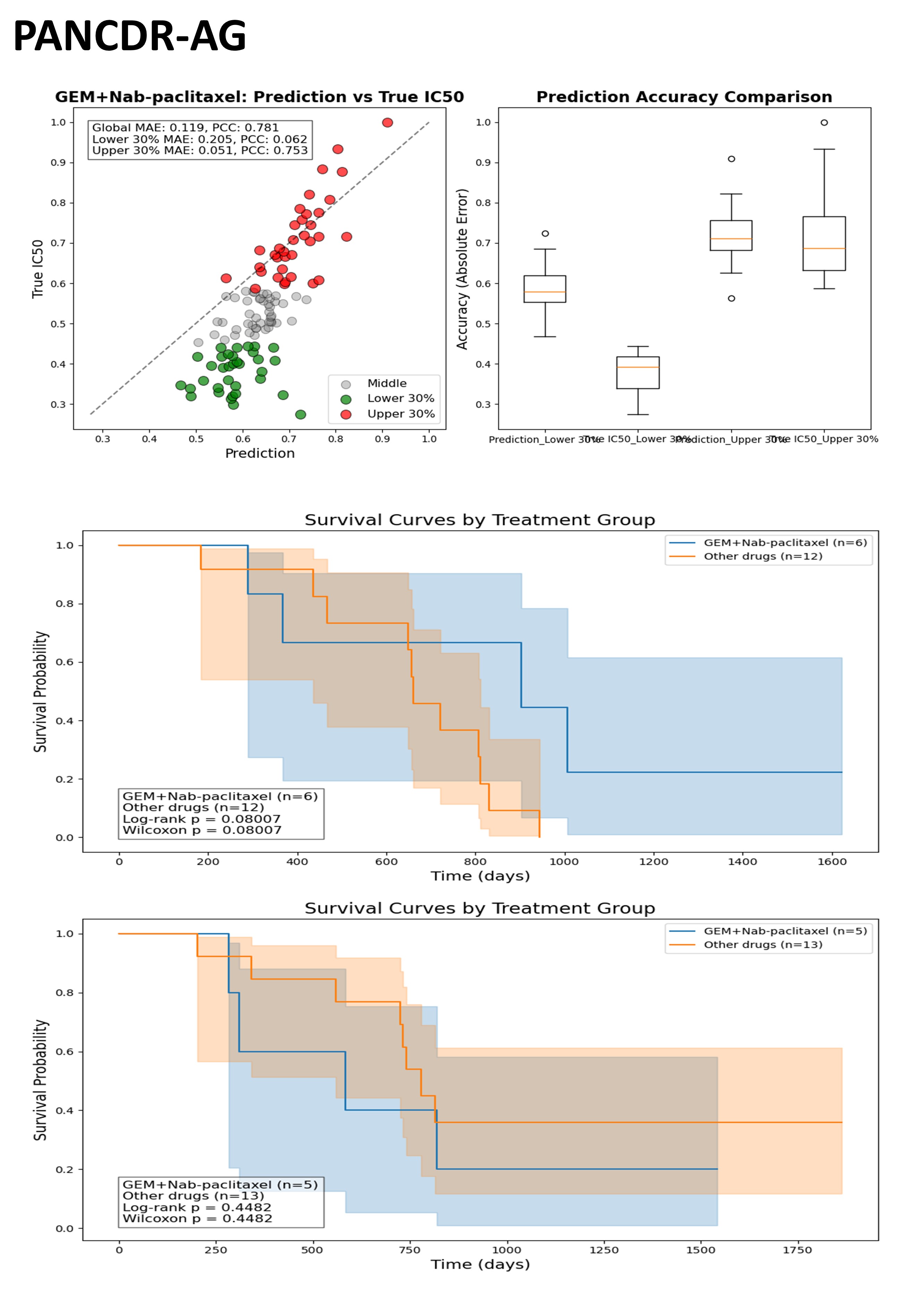

#### Supplementary Figure 11: Performance of PANCDR-guided AG Personalized Treatment Recommendations Im-proves Patient Overall Survival.

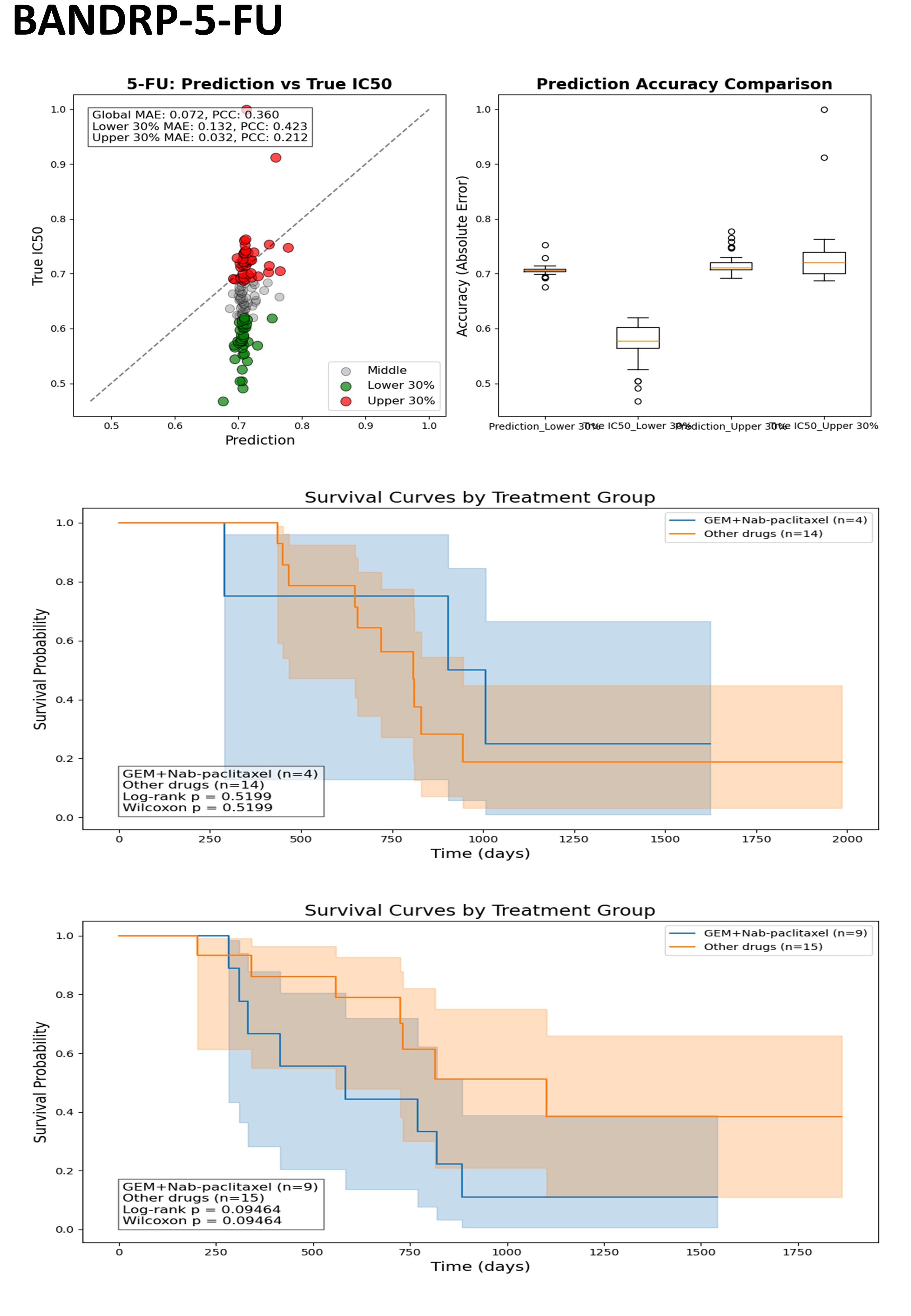

#### Supplementary Figure 12: Performance of BANDRP-guided 5-FU Personalized Treatment Recommendations Im-proves Patient Overall Survival.

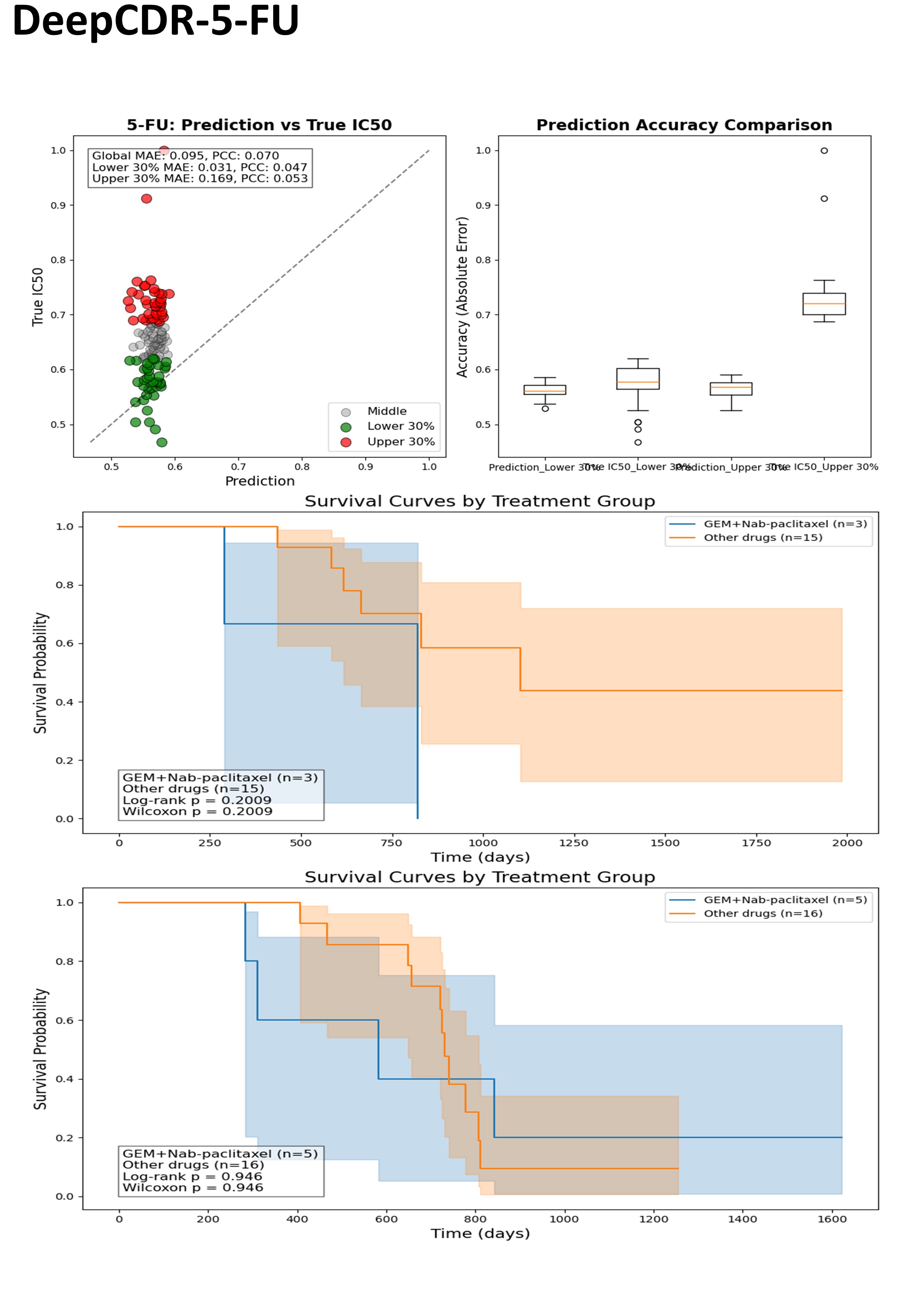

#### Supplementary Figure 13: Performance of DeepCDR-guided 5-FU Personalized Treatment Recommendations Im-proves Patient Overall Survival.

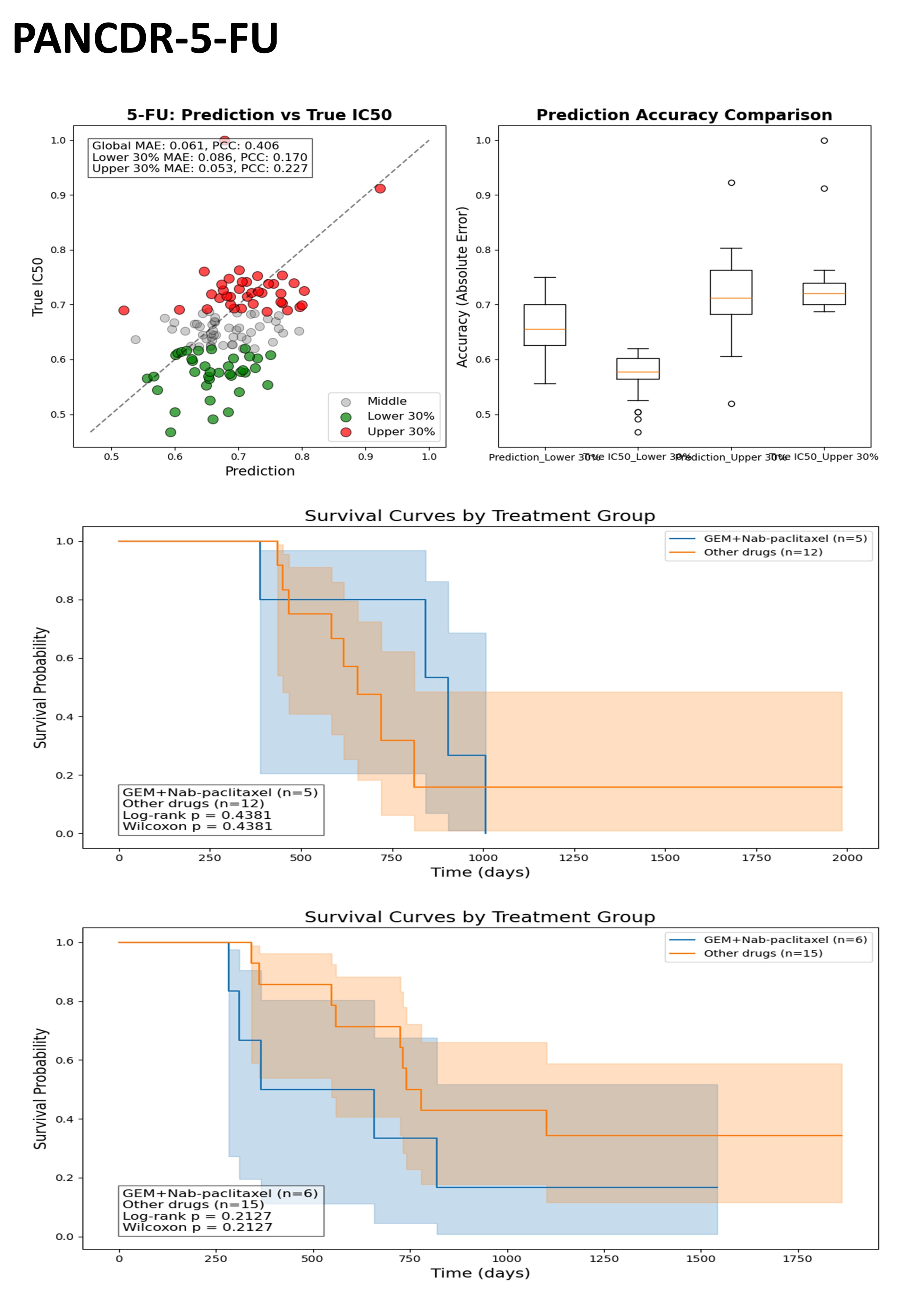

#### Supplementary Figure 14: Performance of PANCDR-guided 5-FU Personalized Treatment Recommendations Im-proves Patient Overall Survival.

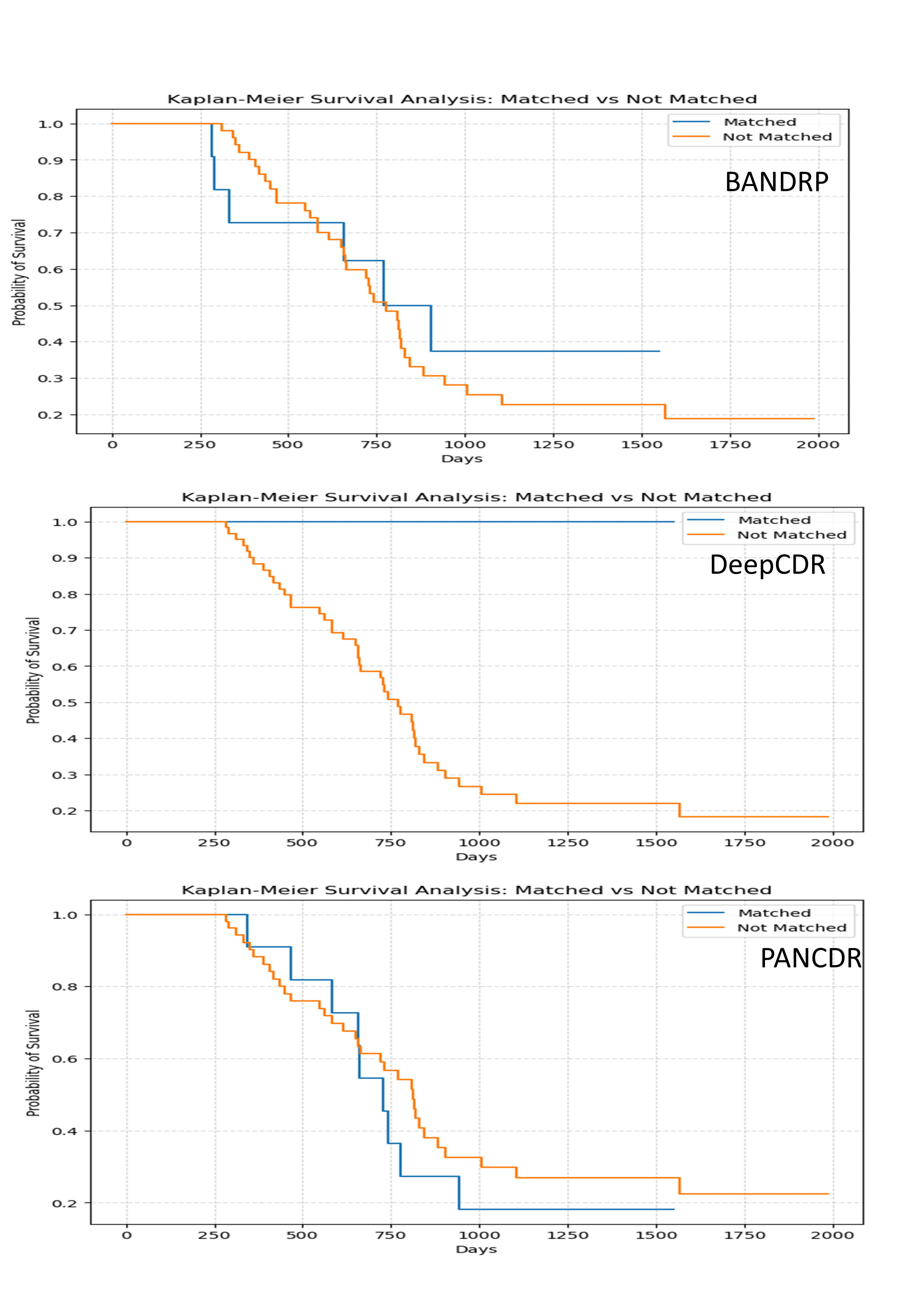

#### Supplementary Figure 15: Performance of methods in the task of ranking the top 8 drug recommendations for each patient based on 76 drugs predicted norm DR-AUC values

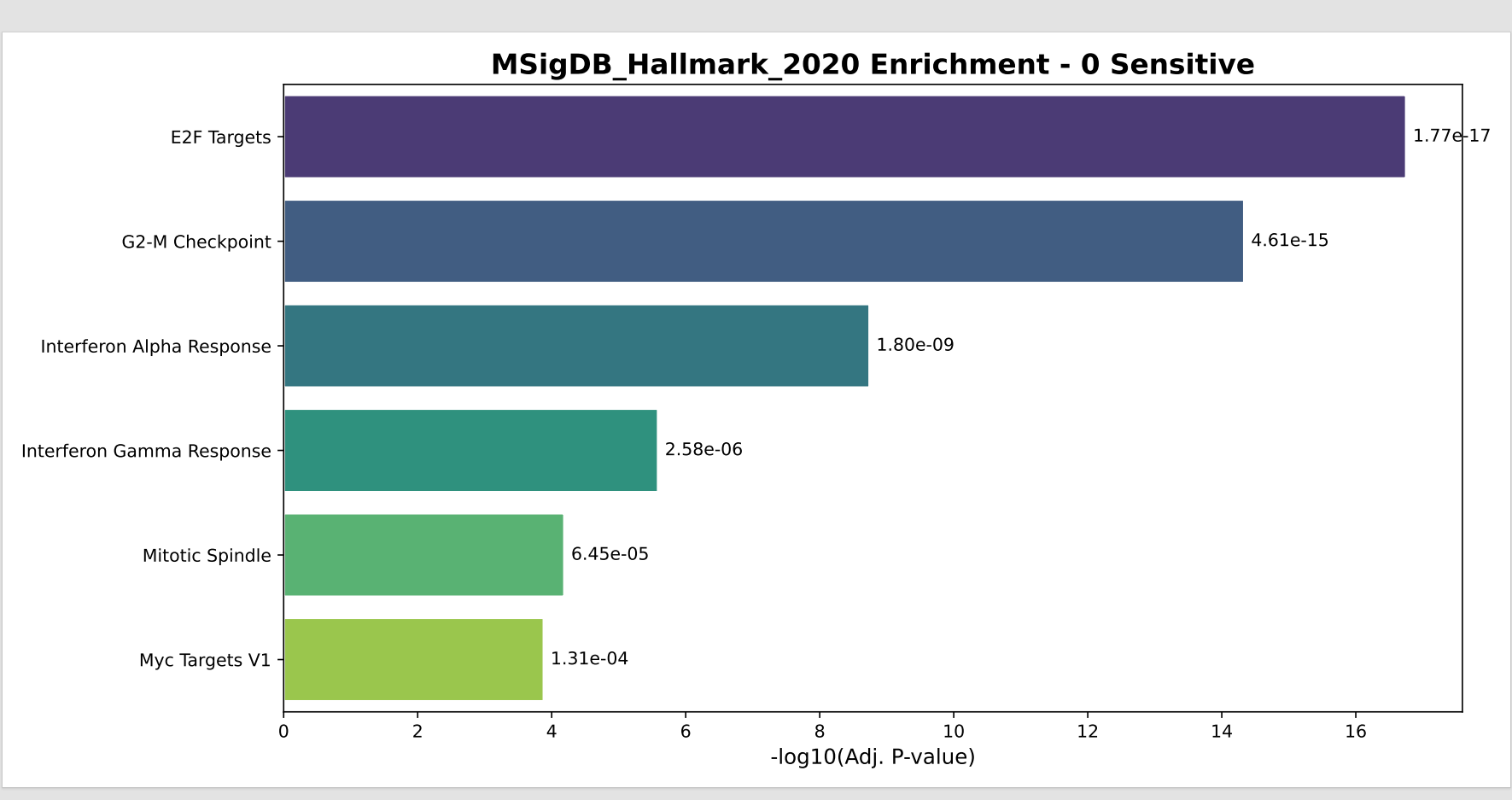

#### Supplementary Figure 16: Hallmark pathway enrichment analysis of transcriptional profiles in Chemo Sen 0 (0). Gene set enrichment analysis (GSEA) was performed using the MSigDB Hallmark 2020 database to identify pathways significantly enriched in the transcriptional profiles of drug-sensitive (0 Sensitive) organoid samples. The top six significantly enriched gene sets are shown, ranked by –log₁₀(adjusted p-value). Cell cycle–related pathways, including E2F Targets and G2-M Checkpoint, exhibited the strongest enrichment, indicating heightened proliferative activity in the sensitive phenotype. Immune-related hallmarks such as Interferon Alpha Response and Interferon Gamma Response were also significantly upregulated, suggesting potential engagement of innate immune signaling. Adjusted p-values were computed using the Benjamini–Hochberg method.

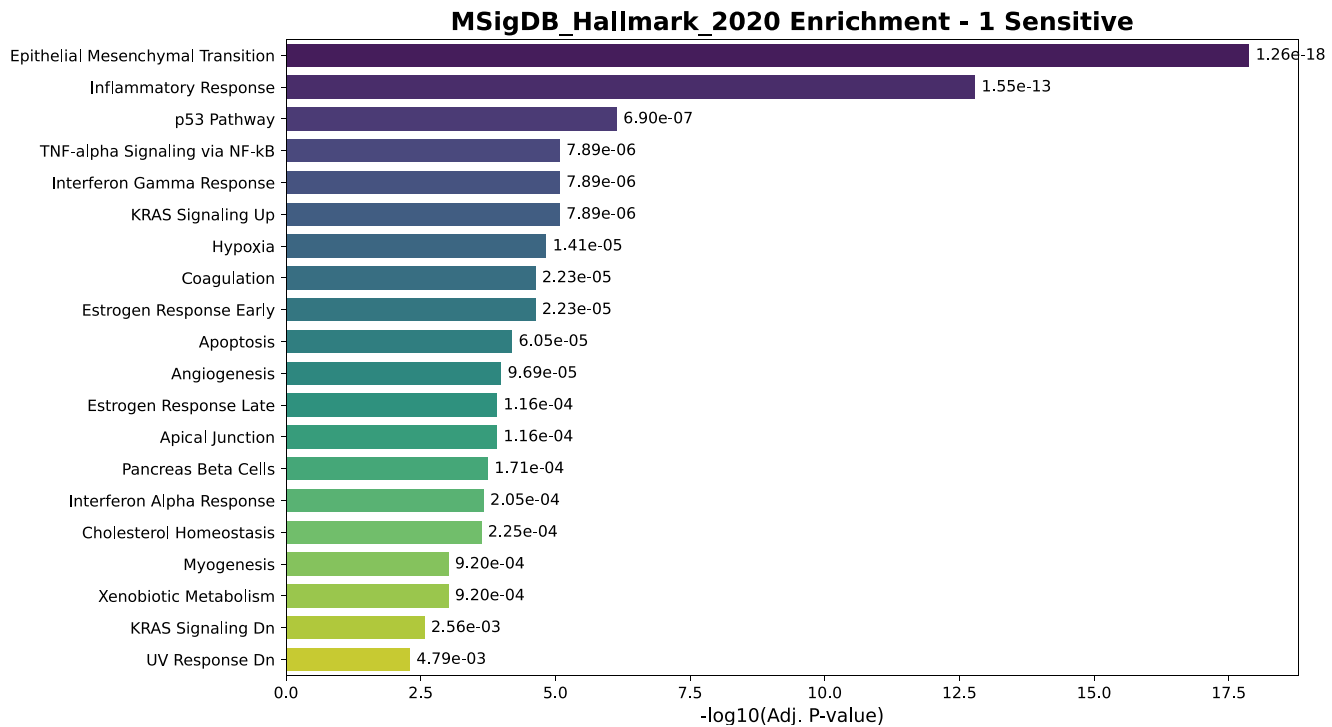

#### Supplementary Figure 17: Hallmark pathway enrichment analysis of transcriptional profiles in Chemo Sen 1 (1). Bar plot showing the top enriched Hallmark gene sets in the drug-sensitive (1 Sensitive) organoid subgroup, based on gene set enrichment analysis (GSEA) using the MSigDB Hallmark 2020 database. Gene sets are ranked by statistical significance (–log₁₀ adjusted p-value, Benjamini–Hochberg correction). The most significantly enriched pathway was Epithelial Mesenchymal Transition (EMT) (adjusted p = 1.26 × 10⁻¹⁸), followed by Inflammatory Response and p53 Pathway, suggesting that sensitivity is associated with mesenchymal features, immune activation, and stress signaling. Additional enrichment of TNF-α and interferon signaling pathways further supports the involvement of inflammatory and apoptotic processes. Metabolic and cell fate–related pathways (e.g., Xenobiotic Metabolism, Cholesterol Homeostasis, Apoptosis) were also significantly represented.

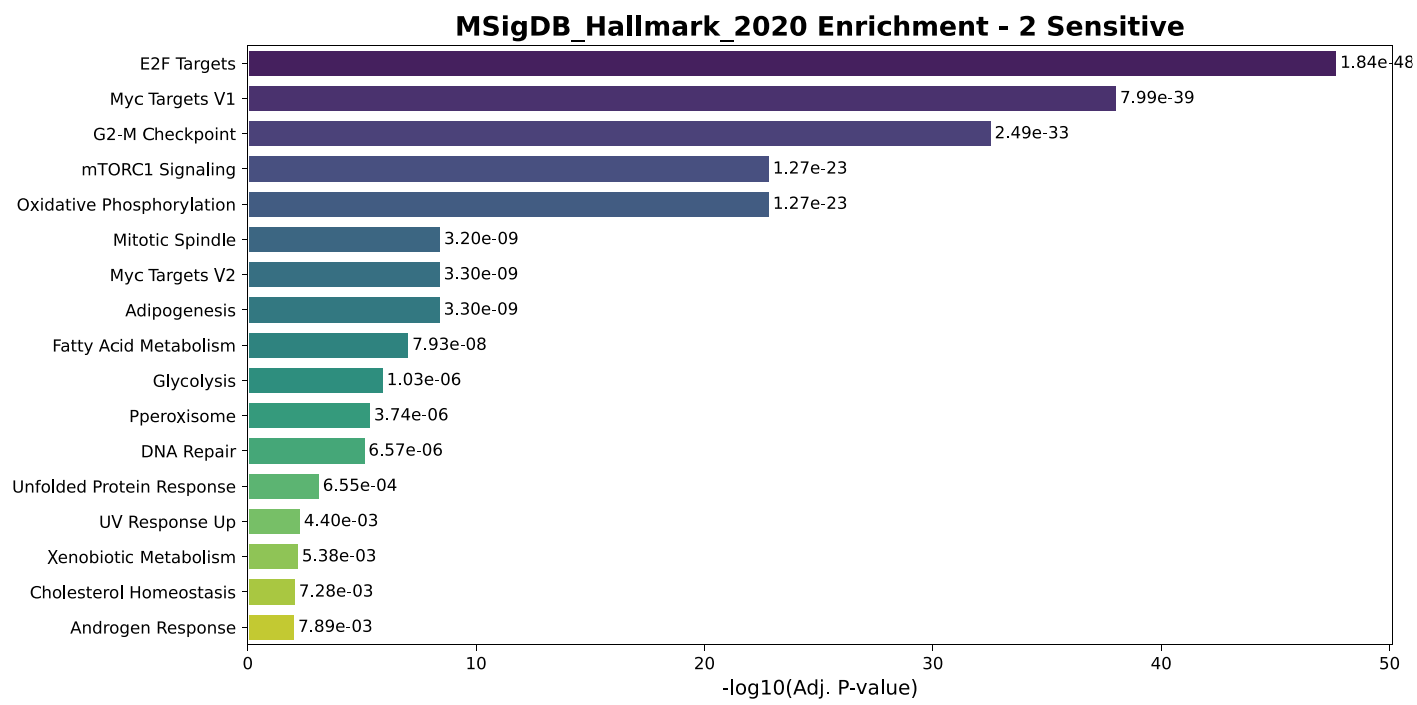

#### Supplementary Figure 18: Hallmark pathway enrichment analysis of transcriptional profiles in Others. Sen 0 (0). Gene set enrichment analysis (GSEA) of the 2 Sensitive subgroup using the MSigDB Hallmark 2020 database. The bar plot shows the top enriched pathways ranked by –log₁₀(adjusted p-value), with adjusted p-values calculated via Benjamini–Hochberg correction. Strong enrichment was observed for cell cycle–associated pathways, including E2F Targets (adjusted p = 1.84 × 10⁻⁴⁸), Myc Targets V1 (adjusted p = 7.99 × 10⁻³⁹), and G2-M Checkpoint (adjusted p = 2.49 × 10⁻³³), suggesting hyperproliferative transcriptional programs in this subgroup. Metabolic pathways such as mTORC1 Signaling, Oxidative Phosphorylation, and Fatty Acid Metabolism were also significantly enriched, indicating a potential coupling of proliferative and bioenergetic processes. Additional pathways, including DNA Repair and Unfolded Protein Response, may reflect stress adaptation mechanisms associated with drug sensitivity.

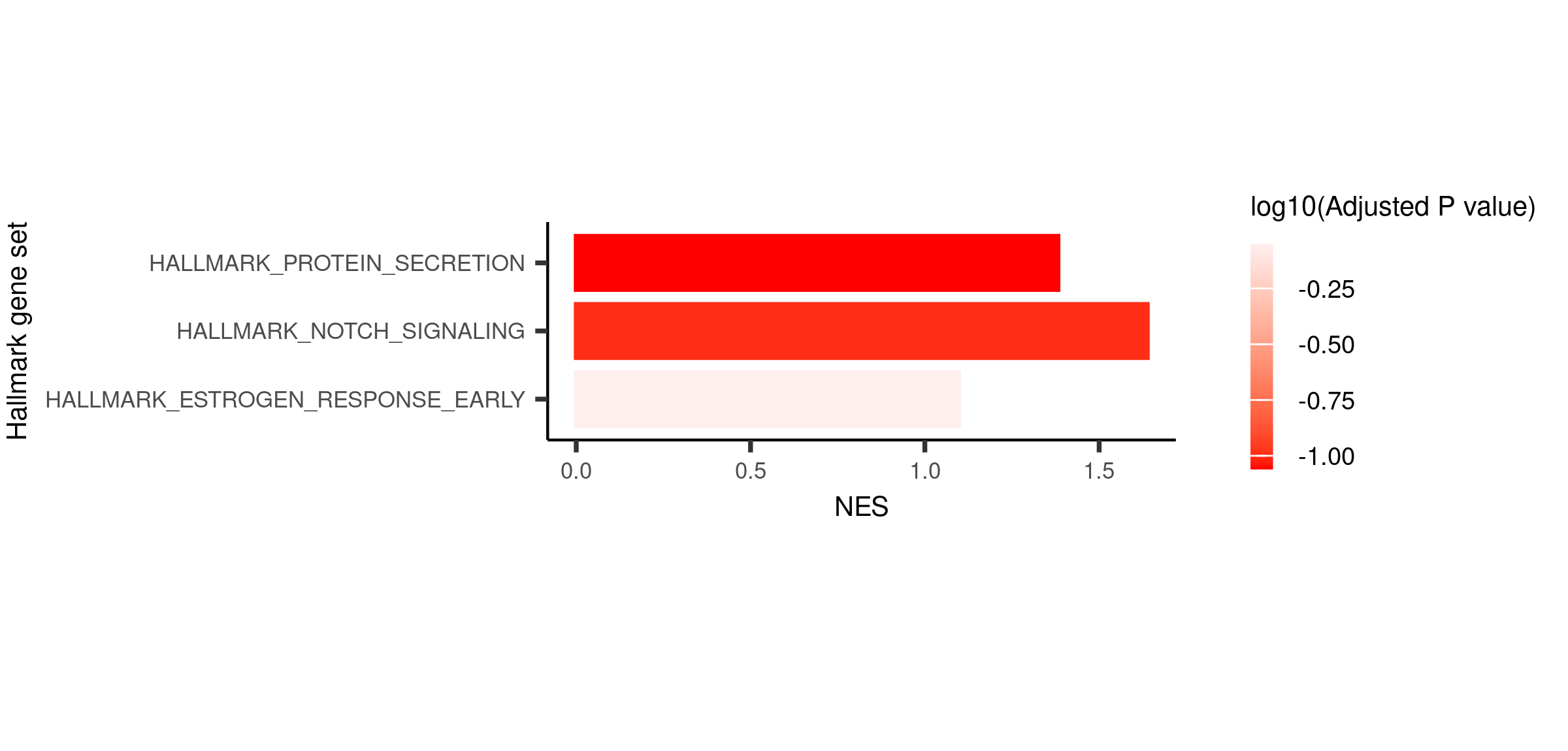

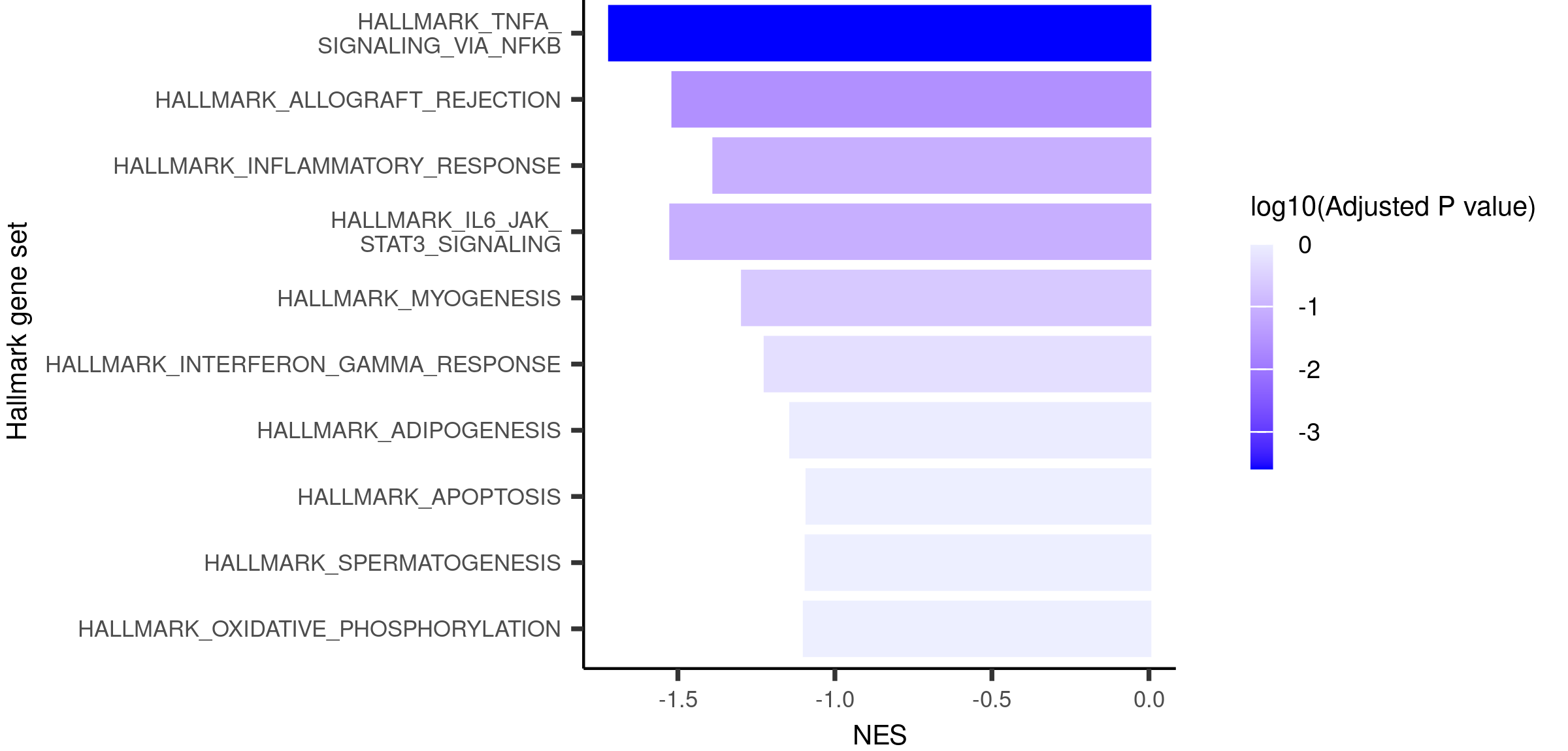

#### Supplementary Figure 19: Contrasting Hallmark pathway enrichment in drug-sensitive versus drug-resistant organoid subgroups.

(a) Top positively enriched Hallmark gene sets in the drug-sensitive subgroup, identified by gene set enrichment analysis (GSEA) using the MSigDB Hallmark 2020 database. The most significantly enriched pathways included Notch Signaling, Protein Secretion, and Estrogen Response (Early), with normalized enrichment scores (NES) > 1.0 and adjusted p-values < 0.1 (Benjamini–Hochberg correction). These pathways suggest active intercellular communication, secretory activity, and hormonal responsiveness may be features of drug-sensitive phenotypes.

(b) Top negatively enriched Hallmark gene sets in the drug-sensitive subgroup (i.e., enriched in the resistant group). Strong negative enrichment was observed for inflammatory and immune signaling pathways, including TNF-α Signaling via NF-κB, Allograft Rejection, and IL6-JAK-STAT3 Signaling, along with Apoptosis and Oxidative Phosphorylation. The enrichment of these pathways in the resistant group may reflect elevated pro-inflammatory signaling, stress responses, and metabolic reprogramming associated with drug resistance.

Color bars indicate –log₁₀(adjusted p-value), with darker shades corresponding to higher statistical significance. Together, these results reveal distinct transcriptional programs underlying drug sensitivity and resistance in organoid models.

### Supplementary Note

#### Note 1

To facilitate biologically meaningful stratification of continuous predictors, we applied a survival-adapted labeling approach based on maximally selected rank statistics. Specifically, we employed the surv_cutpoint function (survminer R package) to determine the optimal threshold of the variable Sensitive in relation to overall survival. This method systematically searches for a cutpoint that best separates the population into two groups with distinct survival outcomes, as determined by maximizing the standardized log-rank statistic. Using overall survival length (OS_length) as the time variable and survival status (Os_status) as the event indicator, the algorithm identified a data-driven cutoff that dichotomizes the Sensitive values into “low” and “high” subgroups. This binarization captures the prognostic inflection point of the variable with respect to survival. The result was visualized using the associated plot() function, which displays the distribution of values and the selected cutpoint, with the two subgroups represented by distinct colors (light blue and red). This survival-informed labeling procedure enables consistent downstream analyses, ensuring that group-level comparisons reflect outcome-relevant biological stratification.
